# “Epidermal Eg5 promotes X-ROS dependent paclitaxel neurotoxicity”

**DOI:** 10.1101/2023.05.22.541784

**Authors:** Antonio Cadiz Diaz, Anthony M Cirrincione, Natalie A Schmidt, Marie J Ugo, Maria Celina Amaya Sanchez, Cassandra A. Reimonn, Stefan Wuchty, Adriana D Pellegrini, Leah RK Rude, Leah G Pappalardo, Daniel P Regan, Caitlin Howell, Sybil Hrstka, Surendra Dasari, Thomas S Lisse, Benjamin J Harrison, Mike Xiangxi Xu, Nathan P Staff, Sandra Rieger

## Abstract

Taxanes are chemotherapeutic agents that induce microtubule modifications in cancer cells, resulting in cell cycle modifications and tumor remission. Here we show that paclitaxel, a widely used taxane, also induces microtubule modifications in healthy epidermal keratinocytes leading to chemotherapy-induced peripheral neuropathy (CIPN). Paclitaxel activates the cell cycle regulator, Kinesin-5 (Eg5), which promotes microtubule detyrosination and fasciculation (dfMT). Eg5 loss protects neurons from paclitaxel neurotoxicity, whereas keratinocyte-specific overexpression promotes axon degeneration. *In vivo* imaging and 3D reconstructions of dfMTs and nuclei, combined with mechanotransduction studies further show that dfMTs constrict keratinocyte nuclei, leading to nuclear Nox-dependent reactive oxygen species (X-ROS) formation upstream of MMP-13 and cutaneous sensory axon degeneration. This new insight facilitates our understanding of chemotherapy side effects and highlights the need for targeted therapies.

## Introduction

A wide variety of chemotherapeutic agents, including paclitaxel (Brand name Taxol), cause sensory-dominant peripheral neuropathy^1-8^. Studies have shown that the earliest signs of degeneration are detected in the unmyelinated sensory neuron population that innervates the epidermis^9^. Symptoms can range from pain and tingling to temperature sensitivity and numbness originating in the hands and feet and progressing proximally^10^. One in every two chemotherapy patients suffers from chemotherapy-induced peripheral neuropathy (CIPN), and one in every three patients requires a dose reduction or discontinuation of this life-saving treatment^11^, therefore decreasing the chance of survival. There are currently no treatments available to prevent or reverse CIPN. Recent research, including our own, has focused on the molecular mechanisms by which paclitaxel damages healthy cells. Putative mechanisms for paclitaxel neurotoxicity have been suggested based on rodent *in vitro* and *in vivo* studies^10^, including reduced local mRNA translation due to decreased axonal microtubule transport and changes in mitochondrial functions^12,13^, leading to the deregulation of intracellular calcium dynamics and neuropeptide release in sensory neurons^14^. Also the induction of inflammatory cascades in rat dorsal root ganglion neurons following paclitaxel treatment involving chemokines, such as CXCL1/8/ MCP-1/CCL-2 and their cognate receptors has been described^15^. Our own studies in zebrafish further discovered that paclitaxel initially damages epidermal keratinocytes prior to promoting cutaneous sensory axon degeneration^16^. We demonstrated that paclitaxel treatment stimulates the formation of reactive oxygen species (ROS) in epidermal keratinocytes, which induces Matrix-Metalloproteinase 13 (MMP-13, collagenase-3) expression that results in extracellular matrix (ECM) degradation and axon degeneration^16,17^. We further showed that pharmacological inhibition of MMP-13 in zebrafish, rats, and mice rescues paclitaxel neurotoxicity and restores epidermal integrity. Studies in Drosophila and mice subsequently demonstrated that overexpression of the collagen-binding β-integrin 1 (ITGB1) in sensory neurons also rescues paclitaxel neurotoxicity^18^, consistent with a model in which paclitaxel induces MMP13 expression in a ROS-dependent manner, leading to epidermal collagen degradation and neuronal downregulation of integrin receptors. This in turn promotes axonal detachment from the ECM and axon degeneration.

Here we investigated the mechanisms by which paclitaxel stimulates ROS formation in epidermal keratinocytes upstream of MMP-13. Because paclitaxel stabilizes microtubules, we reasoned that altered microtubule functions may play a role. Microtubules undergo a process of dynamic instability during which tubulin dimers polymerize and depolymerize via guanosine triphosphate (GTP)/guanosine diphosphate (GDP) cycling whereby bound GTP hydrolyses to GDP during microtubule catastrophe^19^. Paclitaxel inhibits this process and thereby stabilizes the microtubule lattice^20^. Studies have shown that stabilized microtubules are post-translationally modified, or detyrosinated, via removal of the terminal tyrosine on α-tubulin by tubulin carboxypeptidase^21^, thereby exposing a glutamate residue^22^. The regulation of microtubule detyrosination (dMT) and the functional consequence in pathologies is poorly understood. Experiments in cardiomyocytes suggest that dMT formation decreases cell elasticity, and this may underlie heart disease^23,24^. Other studies have associated dMTs with a poor prognosis in cancers, such as breast cancer^25^, and linked them to neurological conditions like Alzheimer’s disease^26^. Microtubules are known mechanotransducers that generate mechanical forces to create instructive signals for cell biological processes, including the regulation of cell polarity^27^, division^28^, and fate^29^. Chronic microtubule stabilization may inadvertently alter cellular signaling processes and promote pathological conditions. Altered mechanical tension in myocytes, for example, has been shown to activate Nox-dependent ROS formation, termed X-ROS^23,24,30-34^. Here we analysed mechanisms by which paclitaxel induces ROS formation in keratinocytes upstream of its neurotoxic effects, which identified Eg5 as a critical mediator.

## Results

### Paclitaxel treatment promotes microtubule detyrosination and fasciculation in epidermal keratinocytes

We first set out to determine whether paclitaxel treatment of larval zebrafish stimulates microtubule stabilisation. Stabilised microtubules undergo a modification whereby the terminal tyrosine residue is removed, leading to detyrosination (dMT)^35^, which is associated with particularly long-term stabilized microtubule populations^36^. Exposure of the penultimate glutamate residue can be detected using a GluTub antibody. Based on our previous findings showing that cutaneous axon degeneration is most prevalent in the distal caudal fin of zebrafish, which consists of two layers of epidermal keratinocytes that are infolded. Each of the two layers are innervated by unmyelinated axons of somatosensory neurons and mesenchymal cells are medially located between the infolded epidermis (**Fig. S1a,b**). This distal fin axon degeneration model was used in all subsequent studies. Caudal fin keratinocytes in vehicle (0.05% DMSO)-treated animals did not show stable microtubule populations following treatment for 3 and 96 hours (**Fig. 1a-c**). However, dMTs were visible in mesenchymal cells at 2 days post fertilization and could be distinguished by their elongate shape spanning multiple keratinocytes (**Fig. 1c, S1a-c, Movie 1**). Paclitaxel (22μM) treatment significantly promoted microtubule detyrosination in keratinocytes following long-term (96hr) but not short-term (3hr) treatment (normalized fluorescence intensity ratio: *3hr (2dpf), vehicle*: 1.039±0.093 vs. *paclitaxel*: 1.197±0.053; *96hr (6dpf), vehicle*: 1.203±0.115 vs. *paclitaxel*: 2.621±0.14)(**Fig. 1b**). Intriguingly, dMTs became fasciculated in a subset of keratinocytes and looped around the cell periphery, most prominently in the distal caudal fin in which axon degeneration commences^16^ (**Fig. 1c, S1d, Movies S2,3**). Microtubule detyrosination, fasciculation, and looping (abbreviated as dfMT) was evident after 48hr paclitaxel treatment in ∼45% of analysed animals, and in ∼90% after 96 hours, but never in vehicle controls, or after 3hr paclitaxel treatment (**Fig. 1d, e**). dfMT presence ranged from 0 to ∼20 keratinocytes/100μm^2^/animal but on average, these were detected in 3.64±1.28 keratinocytes/100μm^2^ following 48hr paclitaxel treatment, and in 7.7±1.6 keratinocytes/100μm^2^ following 96hr paclitaxel treatment (**Fig. 1f**). Also a low dose of paclitaxel (100nM) promoted dfMT formation, but the overall number of affected keratinocytes per animal was significantly lower compared with the high dose (*100nM paclitaxel*: 0.4±0.21 dfMT-positive keratinocytes/100μm^2^)(**Fig. 1f**), suggesting that the extent of dfMT formation is dose-dependent. Interestingly, despite the low number of keratinocytes harbouring dfMTs with 100nM paclitaxel, the fascicle width of dMTs was significantly enhanced (*100nM paclitaxel*: 6.0±0.49μm vs. *22μM paclitaxel*: 3.4±0.17μm)(**Fig. 1g**). To determine if this phenotype was inherent to zebrafish or conserved in mammals, we analysed dfMT formation in the mouse epidermis. A single injection of either vehicle or 20mg/kg paclitaxel into mice (5-6 weeks) resulted in dfMT formation in a subset of suprabasal keratinocytes. These occasionally formed large cell clusters with rosette-like dfMTs connecting them (**Fig. 1h**). In contrast, vehicle-injected control animals never showed dfMT formation although some degree of detyrosination was present.

**Figure 1.**
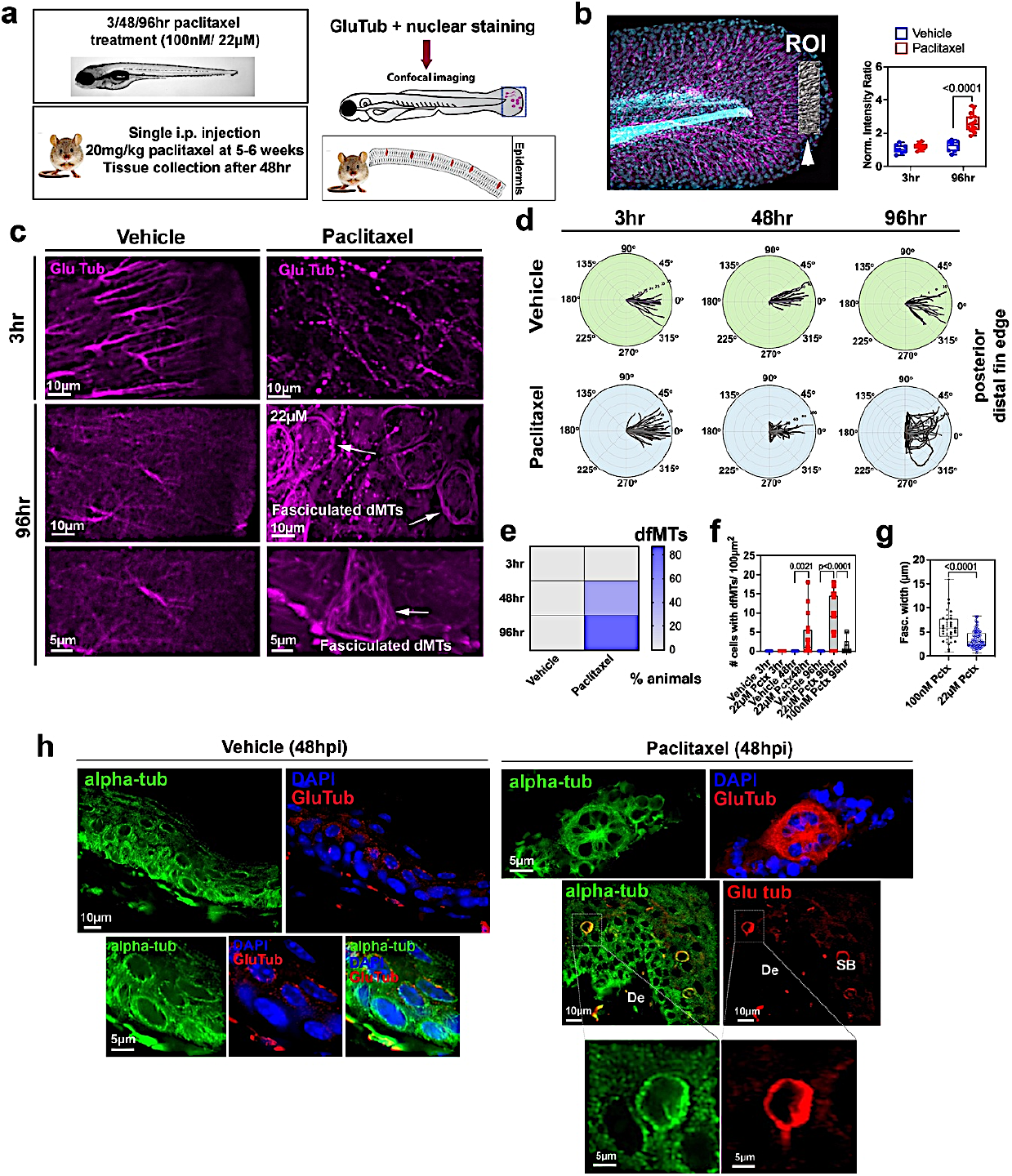
Detyrosination and fasciculation of microtubules following paclitaxel treatment. (**a**) Experimental design for GluTub staining. (**b**) Normalized intensity ratios of GluTub stained microtubules in the caudal fin (see region of interest, ROI, arrowhead) shows significantly increased detyrosination following 96hr, but not 3hr, paclitaxel treatment compared with vehicle (0.05% DMSO) controls. GluTub staining was normalized to Hoechst33342 nuclear stain within each fin. (**c**) Strong GluTub labelling in linear mesenchymal cells at 2dpf following 3hr vehicle treatment (see **Movie S1**), which is absent in these cells in the presence of paclitaxel but instead axons are labelled. Diffuse dMT staining is present in the fin following 96hr vehicle treatment, whereas 96hr paclitaxel treatment shows dfMTs in keratinocytes (arrows, **Movies S2,3**). **(d)** Polar plots depicting orientation and shape of dMTs in the caudal fin (180°=proximal; 0°= distal fin) (n=4 animals/plot). **(e)** Significant increase in the percent animals with dfMTs after 48 and 96hr paclitaxel but not vehicle treatment (n≥15 animals/group). (**f**) Significant increase in the number of caudal fin keratinocytes per 100μm^2^ with dfMTs following 22μM paclitaxel treatment, while few keratinocytes harbour dfMTs when animals are treated with 100mM paclitaxel (n≥17 animals/group). (**g**) dfMTs in animals treated with 100nM paclitaxel for 96hr show increased fasciculation widths compared with 22μM paclitaxel treatment (n=5 animals/group). (**h**) Single injection of 20mg/kg paclitaxel into mice at 5 weeks causes dfMT and rosette formation 48hr after the injection, assessed with alpha-tubulin staining. dfMT formation is evident in the suprabasal epidermis.

Together, these findings demonstrate that paclitaxel treatment promotes a dose-dependent dfMT formation in subpopulations of epidermal keratinocytes, which may contribute to H_2_O_2_ dependent MMP-13 expression and neurotoxicity.

### Paclitaxel promotes X-ROS formation via altered microtubule mechanotransduction

We next asked whether keratinocyte dfMT formation contributes to ROS production. ROS can be derived from two major sources within cells, mitochondria and membrane-bound NADPH oxidases. We hypothesized that keratinocyte-specific mitochondrial ROS (mitoROS) might be a source because we previously detected morphological changes in keratinocyte mitochondria following paclitaxel treatment^17^, and mitochondrial diseases have been linked to peripheral neuropathy^37-39^. To assess mitoROS levels in keratinocytes, we expressed *tp63*:HyPer-mito, a mitochondria-targeted genetic sensor for the ROS, hydrogen peroxide (H_2_O_2_)^40,41^. Treatment with paclitaxel for 3hr and 48hr, which is when cytoplasmic ROS are abundant^16,42^ did not significantly increase mitochondrial HyPer oxidation (3hr vehicle: 1.054±0.031 vs. paclitaxel: 1.252±0.066; 48hr vehicle: 1.029±0.086 vs. paclitaxel: 0.972±0.2)(**Fig. S2**). This suggested that paclitaxel is not contributing to mitoROS formation and that NADPH oxidases are the likely source of ROS in keratinocytes, consistent with our previous findings that 3hr and 48hr paclitaxel treatment induces cytoplasmic H_2_O_2_ production^16,17^.

NADPH oxidases were therefore an alternative source for ROS. The NADPH oxidase family consists of Nox1-5 and Duox1/2, with Duox2 being species-dependent^43^ and Nox1-4 being activated by the subunit p22phox^44,45^. We first used quantitative PCR (qPCR) in zebrafish to analyse the expression of the two known epithelium-specific NADPH oxidases, *nox1* and *duox*, in addition to *nox2*, which was shown to be activated by mechanotransduction mechanisms in cardiomyocytes^32^. QPCR on whole larval zebrafish showed only *nox1* being significantly upregulated following 48hr paclitaxel treatment (fold-change from vehicle control: *duox*: 1.63±0.23, *nox1*: 2.5±0.61, *nox2*: 1.35±0.12)(**Fig. 2a**). Immunofluorescence staining and 3D reconstruction of Nox1, and nuclear staining using Hoechst33342, further showed a time-dependent recruitment of Nox1 to the plasma membrane and nucleus in epidermal keratinocytes following paclitaxel, but not vehicle, treatment (membrane:cytoplasmic ratio, 48hr *vehicle*: 1.23±0. 04 vs. *paclitaxel*: 1.5±0.09; *120hr vehicle:* 1.74±0.09 vs. *paclitaxel*: 2.63±0.29)(**Fig. 2b**)(**Movies S4,5**). These findings indicate that paclitaxel modulates Nox1 activity by promoting its subcellular translocation, which could be mediated by the altered mechanotransduction of dfMTs, consistent with findings in muscle cells.

**Figure 2.**
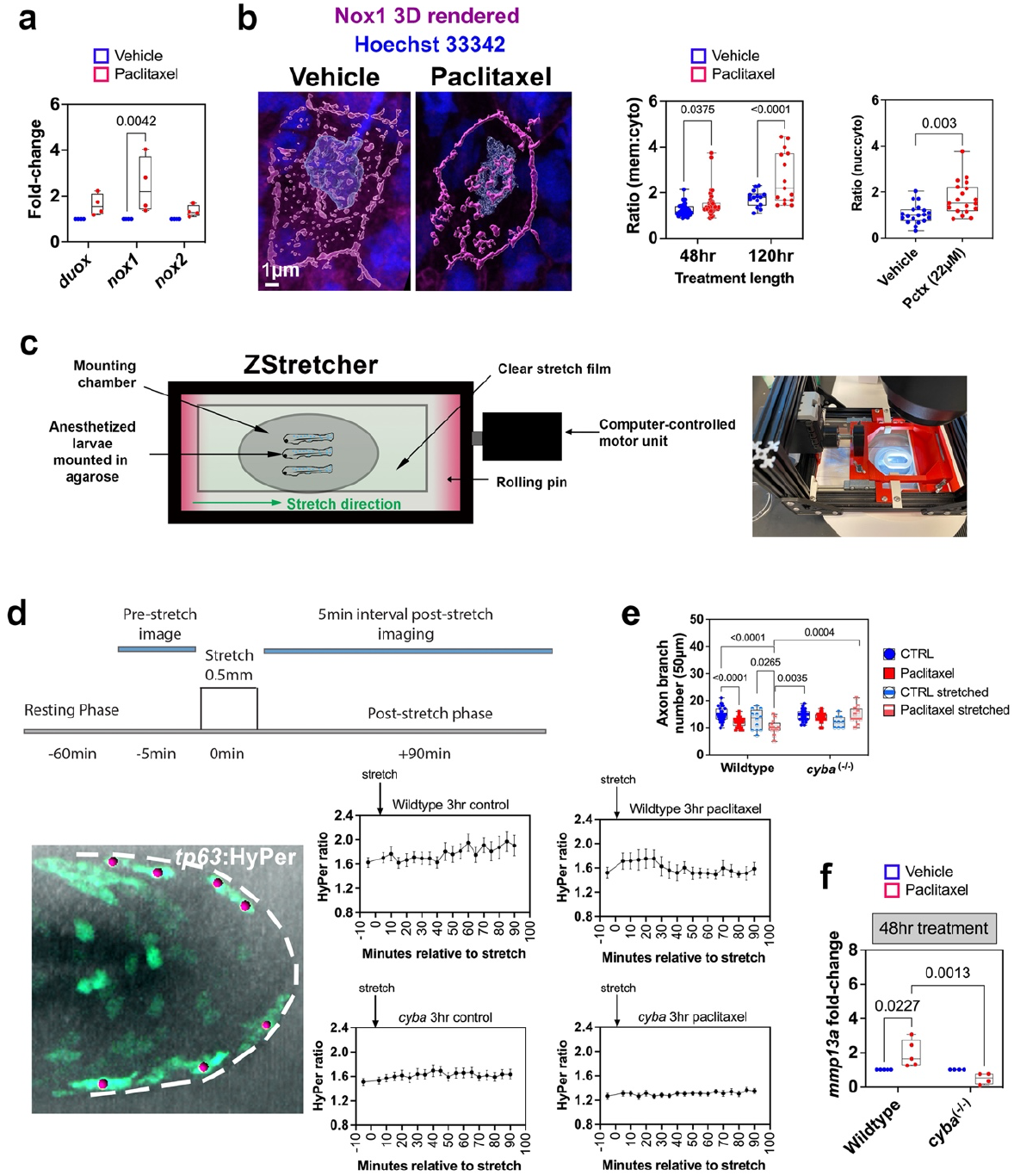
Mechanotransduction activates Nox1 upstream of MMP-13 and cutaneous axon degeneration. (**a**) Quantitative PCR (qPCR) shows significantly increased *nox1* expression in 4dpf zebrafish following 48hr paclitaxel treatment (i.e. axon degeneration onset)(n=3 biological replicates with 10 animals per replicate). (**b**) Nox1 is distributed in the cytoplasm and nucleus of caudal fin keratinocytes when fish are treated with vehicle (0.05% DMSO) and translocates to the plasma membrane and clusters in the nucleus (arrow) following prolonged paclitaxel treatment (see **Movies 4,5**). Membrane:cytoplasmic ratio of Nox1 staining increases upon continued paclitaxel treatment (n≥6 animals/group), similar to the nuclear:cytoplasmic ratio (n≥9 animals/group). (**c**) ZStretcher design for simultaneous stretching and confocal imaging of live zebrafish. (**d**) Ratiometric HyPer imaging from 5min pre whereas HyPer oxidation is largely reduced or absent in vehicle and paclitaxel-treated *cyba*^*-/-*^ mutants, respectively (n=5 animals/group). (**e**) Prolonged paclitaxel, but not vehicle, treatment reduces the number of axon branches in the caudal fin of wildtype but not *cyba*^*-/-*^ fish (n=6-9 animals/group). Stretch slightly enhances axon degeneration in wildtype fish treated with paclitaxel, but not *cyba*^*-/-*^ fish. (**f**) qPCR demonstrates enhanced *mmp13a* (MMP13 homolog) expression in wildtype fish treated with paclitaxel, but not in *cyba*^*-/-*^ mutants (n=10-20 animals/group in 3 biological replicates).-stretch to 90min post-stretch in wildtype and *cyba*^*-/-*^ mutants treated either with vehicle or paclitaxel demonstrates faster activation dynamics for H_2_O_2_ production in wildtype animals treated with paclitaxel, with paclitaxel, whereas HyPer oxidation is largely reduced or absent in vehicle and paclitaxel-treated *cyba*^-/-^ mutants, respectively (n=5 animals/group). (e) Prolonged paclitaxel, but not vehicle, treatment reduces the number of axon branches in the caudal fin of wildtype but not *cyba*^-/-^ fish (n=6-9 animals/group). Stretch slightly enhances axon degeneration in wildtype fish treated with paclitaxel, but not *cyba*^-/-^ fish. (f) qPCR demonstrates enhanced mmp13a (MMP13 homolog) expression in wildtype fish treated with paclitaxel, but not in *cyba*^-/-^ mutants (n=10-20 animals/group in 3 biological replicates).

To test the hypothesis that paclitaxel induces Nox-dependent ROS formation by altering microtubule tension, we engineered a device (ZStretcher) that allowed us to exert tension on the zebrafish caudal fin via mechanical stretch while simultaneously being able to monitor ROS formation using time-lapse imaging (**Fig. 2c**). We reasoned that stretch in the presence and absence of paclitaxel modifies microtubule dynamics and this should impact ROS production in wildtype fish whereas Nox-deficient *cyba*^*-/-*^ mutants do not produce ROS in response to stretch. The ZStretcher was designed to fit into the stage of an LSM880 confocal microscope (Zeiss) for imaging of live larval zebrafish mounted in agarose on a clear plastic film that can be stretched in 1mm increments using a software-controlled motor unit. First, we characterised the extent of stretch that was induced in wildtype transgenic Tg(*tp63*:GFP-CAAX)^16^ fish in which keratinocyte plasma membranes are fluorescently labelled. Comparison of vehicle-treated pre-and 15min post-stretch keratinocytes showed a significant increase in keratinocyte length, but not width, at the fin edge (keratinocyte length, *medial fin:* pre-stretch: 27.99±0.97μm vs. post-stretch: 30.37±1.05μm; *fin edge:* pre-stretch: 30.35±1.36μm vs. post-stretch: 36.46±2.04μm; keratinocyte width, *medial fin:* pre-stretch: 19.19±0.86μm vs. post-stretch: 19.01±0.69 μm, *fin edge:* pre-stretch: 11.62±0.59 μm vs. post-stretch: 13.52±1.16μm)(**Fig. S3a**). Subsequent fixation of stretched wildtype and *cyba*^*-/-*^ mutant fish followed by immunofluorescence staining for alpha-tubulin confirmed stretching of microtubules in fin edge keratinocytes of vehicle-treated, but not in paclitaxel-treated, animals (**Fig. S3b**), confirming altered mechanical characteristics of stabilised microtubules.

To further investigate whether paclitaxel in combination with stretch alters ROS formation, we performed HyPer imaging. Dual channel imaging of oxidized and unoxidized HyPer in 5min intervals for 90 minutes pre- and post-stretch showed that HyPer oxidation in vehicle control fish was relatively sluggish, with a maximum increase at ∼80min post-stretch (*HyPer ratio (505/420) pre- vs. post-stretch (max)*: 1.62 vs. 1.97)(**Fig. 2d**). Paclitaxel-treated animals (3hr treatment), in contrast, showed rapid HyPer oxidation that peaked between 10-20min (*HyPer ratio pre- vs. post-stretch (max)*: 1.49 vs. 1.75) and thereafter declined. As predicted, HyPer oxidation was Nox-dependent since stretching of *cyba*^*-/-*^ mutants with and without paclitaxel treatment resulted in a 10-fold decrease in HyPer oxidation in the vehicle controls, and the absence of a response to stretch in the paclitaxel group (*cyba*^*-/-*^, *vehicle: HyPer ratio pre- vs. post-stretch (max)*: 1.50 vs. 1.69 vs. paclitaxel: 1.25 vs. 1.36). The remaining HyPer oxidation in vehicle-treated *cyba*^*-/-*^ mutants suggests that changes in MT tension may also stimulate other P22phox-independent NADPH oxidases, such as Duox, or stimulate mitochondrial ROS production. This indicates that dfMTs promote X-ROS formation in keratinocytes following paclitaxel treatment.

To further determine whether microtubule stretch enhances paclitaxel-induced axon degeneration in a Nox-dependent manner, we compared the axon branch number in wildtype and *cyba*^*-/-*^ fish treated with vehicle and paclitaxel either stretched or unstretched. Stretching was performed after 96hr treatment, followed by 2hr post-stretch fixation, and staining for detection of acetylated tubulin in axons. Axon branch quantifications showed a significant decrease in axon branches in the caudal fin of wildtype paclitaxel-treated but not vehicle control fish, and no significant change was found within the *cyba*^*-/-*^ group. While stretching slightly enhanced degeneration within wildtype animals, it was significantly rescued in paclitaxel-treated and stretched *cyba*^*-/-*^ mutants (**Fig. 2e**). To determine whether Nox activation occurs upstream of MMP-13, we analysed *mmp13* expression by qPCR in whole larval wildtype and *cyba*^*-/-*^ mutant fish in the absence and presence of paclitaxel. Wildtype but not *cyba*^*-/-*^ fish treated for 48hr with paclitaxel to stimulate axon degeneration displayed increased *mmp13* expression (**Fig. 2f**). Together these data support a model by which paclitaxel-induced microtubule stabilization induces X-ROS formation and MMP-13 expression upstream of cutaneous sensory axon degeneration.

### Paclitaxel treatment stimulates cell cycle gene expression in the skin

We next sought to investigate the molecular mechanisms underlying paclitaxel-induced microtubule fasciculation. Because we previously identified that CIPN dependence on MMP-13 is conserved^16,17^, we analysed our mouse RNAseq dataset in which mice received four intraperitoneal injections of either vehicle or 2mg/kg paclitaxel every other day four times^42^. In this experiment, the skin was harvested on days 4, 7, 11, and 23 for comparisons of gene expression profiles (**Fig. 3a**). Peripheral neuropathy was correlated with these times points using von Frey behavioural testing, which determined that peak neuropathy was present at day 7 (D7). We further determined gene co-expression profiles (log2 fold-change: ≥0.65)(**Fig. 3b**), followed by gProfiler enrichment analysis (**Fig. 3c**). This identified several biological categories during peak neuropathy with the largest clusters being implicated in extracellular matrix organization (e.g. “ECM”, “collagen-containing ECM”, “collagen trimer”) and cell cycle regulation (e.g. “outer kinetochore”, “kinesin complex”, condensed chromosome, centromeric region”, “mitotic spindle”, “microtubules”), consistent with our findings in zebrafish. Interactome analysis using STRING^46^ to predict protein association networks further identified major clusters involved in cell cycle regulation, which contained various kinesins, microtubule and nucleus regulators, as well as one cluster annotated as “disulfide bond”, consistent with a role for oxidative signal activation^47^ (**Fig. 3d**). Given that paclitaxel is a known cell cycle checkpoint regulator^48,49^, we further analysed the expression of the known checkpoint regulators, *Plk1, Cdc20, Ndc80, Bub1c, Dlgap5*, which showed a co-upregulation from D7 onward (**Fig. S4a**). Additional genes in our data set that were significantly upregulated during peak neuropathy and formed a network included the mitosis regulating genes, including *Cdk1, Aurka, Aurkb*, and *Kif11* (**Fig. S4b**). *Kif11*, which encodes Eg5 (also known as Kinesin-5), was especially interesting since 1) members of the Kinesin family are implicated in microtubule regulation and cell cycle progression^50^, 2) the implication of EG5 in the regulation of cytokinesis via its microtubule crosslinking activity^51^, consistent with our microtubule fasciculation phenotype, and 3) Eg5 inhibition in mice with the small molecule inhibitor, monastrol, has been shown to alleviate CIPN caused by bortezomib^52^. Besides *Kif11*, we also found other significantly upregulated *Kif* genes during peak neuropathy: *Kif2c, Kif14, Kif15, Kif20a, Kif20b and Kif23* (**Fig. 3e**). We further analysed the expression profiles of several *Nox* family genes, which showed that *Cyba* and *Cybb*, the latter being specific to immune cells, were significantly upregulated at D23, whereas other genes, such as *Nox1* and *Nox4* were slightly but not significantly increased at varying days (**Fig. 3f**), suggesting a primary regulation via post-translational mechanisms, consistent with previous findings^32,34,53^.

**Figure 3.**
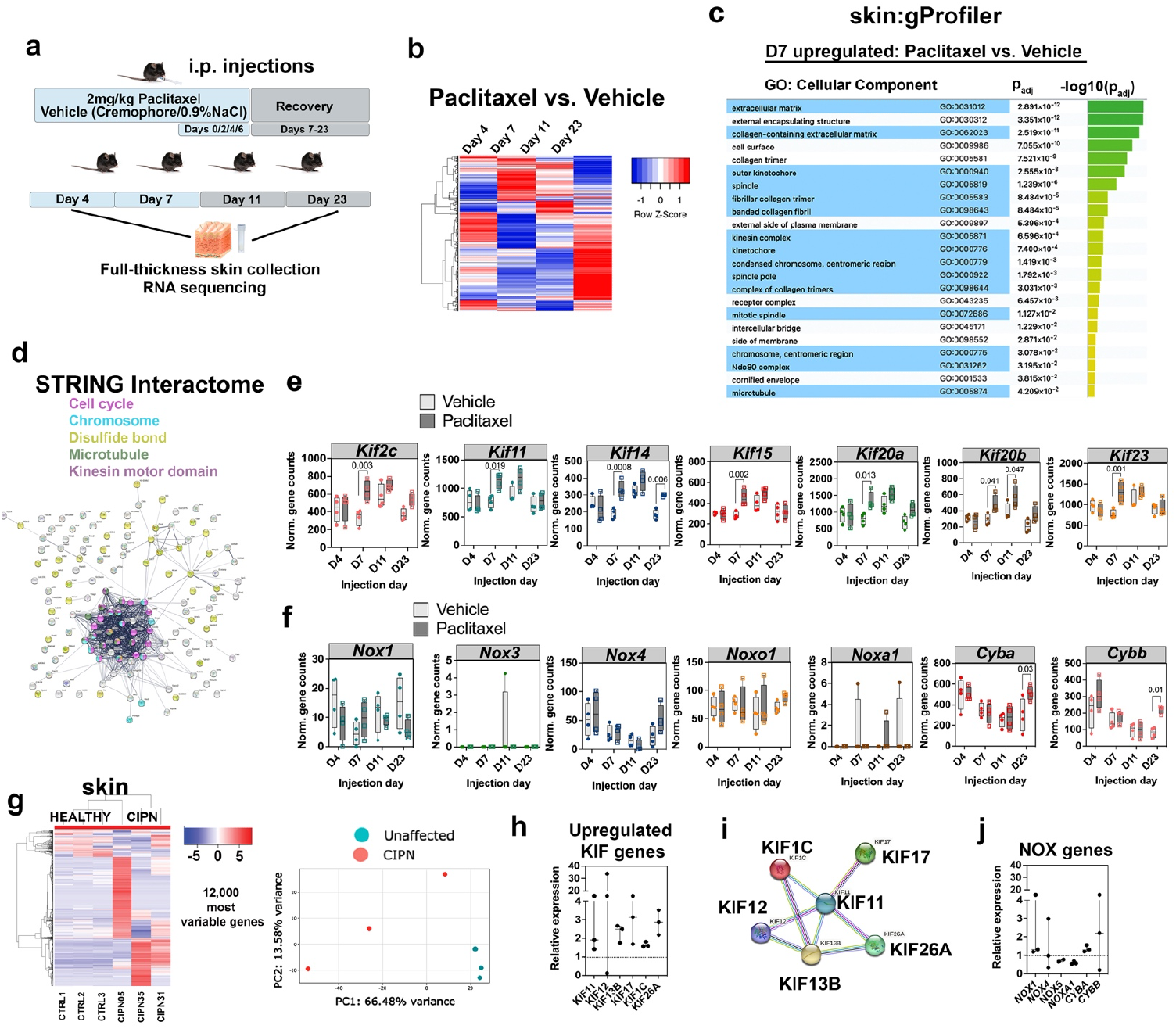
Paclitaxel treatment induces cell cycle genes in the skin of mice and patients with CIPN. (**a-f**) RNAseq of glabrous skin from mouse hind paws. (**a**) Treatment scheme for mouse RNAseq analysis (n=4 animals per group). (**b**) Heatmap showing significantly up (red) and down (blue) regulated genes. (**c**) gProfiler analysis at peak neuropathy day 7 (D7) reveals GO terms for extracellular matrix and collagen-related processes, and cell cycle regulation. (**d**) STRING interactome analysis of genes in (**c**) identifies a major cluster for cell cycle genes (magenta, blue), and proteins involved in disulphide bond formation (yellow). (**e**) Normalized gene counts for significantly upregulated *Kif genes* following paclitaxel treatment. (**f**) Normalized gene counts for Nox genes and their regulators detected in the skin. (**g-j**) RNAseq using human lower leg skin from 3 CIPN patients (CIPN5, 31, 35 weeks) and 3 control subjects (CTRL1-3). (**g**) Heatmap and PCA plot reveals the separation of CIPN patients from the controls. (**h**) Relative gene expression of upregulated KIF genes (left to right: *KIF11, KIF12, KIF13B, KIF17, KIF1C, KIF26A*). (**i**) STRING network analysis reveals KIF11 as a central hub for co-upregulated KIF genes. (**j**) NOX1 is most strongly induced and shows great variability together with NOX4 and CYBB (left to right: *NOX1, NOX4, NOX5, NOXA1, CYBA, CYBB*).

To determine the extent to which these findings are conserved in humans, RNAseq was conducted on three paclitaxel-treated breast cancer patients with CIPN and three age/sex-matched healthy controls. Upon enrolment, each participant completed a questionnaire and physical examination, followed by a full-thickness skin punch biopsy. The patients were diagnosed with CIPN 35 weeks (CIPN01), 31 weeks (CIPN02), and 5 weeks (CIPN03) prior to the biopsy. After RNA extraction and processing, Illumina RNAseq analysis was performed, followed by heatmap and PCA plot generation using iDEP^54^. Differences in gene expression between the three CIPN patients and healthy controls were evident from these two analyses (**Fig. 3g**). Further enrichment analysis showed that the patients with their first CIPN diagnosis 31 and 35 weeks prior to the skin biopsy shared distinct gene clusters that separated from the patient with the most recent CIPN diagnosis (5 weeks prior). Although *KIF11* was upregulated in all three patients compared with the controls, the expression differences were not significant due to a strong expression increase in one of the CIPN patients (CIPN02).

Differential gene expression analysis further identified several other significantly upregulated *KIF* genes (**Fig. 3h**). STRING interactome analysis of upregulated *KIF* genes revealed that KIF11 serves as a central hub for these co-upregulated KIF genes (**Fig. 3i**). Expression analysis of NADPH oxidases identified *NOX1* and CYBB as the most highly (but not significantly) upregulated genes in the paclitaxel-treated patient skin compared with healthy controls (**Fig. 3j**). Using the top 1,200 most variable differentially expressed genes, we further identified three major clusters with iDEP (**Fig. S4c**). The first cluster showed pathways downregulated in the CIPN patients, which included processes, such as “epithelium/epidermis development”, “keratinocyte differentiation”, “gluconeogenesis”, and “lipid catabolic processes”. The second cluster (B) harbored genes upregulated in the 5-week CIPN patient skin, and these were involved in processes like “hydrogen peroxide catabolic process”, “cell death”, and “response to hydrogen peroxide”. Cluster C contained genes upregulated in the 31/35-week CIPN patients, with functions in “chromatin remodeling/assembly/disassembly”, “chromosome condensation”, “chromosome organization”, and “DNA conformation change”. These results establish conserved skin-specific gene expression profiles for mice and humans whereby paclitaxel promotes the expression of genes involved in cell cycle and oxidative stress regulation.

### Kif11/EG5 promotes dfMT and nuclear X-ROS formation in zebrafish

Because Eg5 has been linked to bortezomib-induced peripheral neuropathy, we focused our subsequent analysis on this cell cycle regulator, using primarily zebrafish due to their *in vivo* imaging capabilities. First, we determined whether paclitaxel induces EG5 expression in caudal fin keratinocytes. Following 96hr vehicle treatment, EG5 immunofluorescence staining with an antibody targeting the conserved phosphorylation site, Thr927 (PTGTTPQRK), revealed a uniform punctate Eg5 localization in the caudal fin except around the fin edge (**Fig. 4a**). Treatment with 22μM paclitaxel for 96 hours promoted Eg5 puncta formation and Eg5 location to structures reminiscent of microtubule asters, such as formed during early mitosis^55^, which was most prominent at the fin edge. Because of the microtubule crosslinking activity of Eg5 during spindle formation and elongation^56,57^, which resembles the observed Eg5 expression pattern in keratinocytes, we asked whether Eg5 plays a role in paclitaxel-dependent dfMT formation. Indeed, combination of the Eg5 inhibitor, EMD534085, with 22μM paclitaxel largely prevented dfMTs formation in the majority of animals (*vehicle*: 0%, *EMD534085*: 0%, *100nM paclitaxel:* 20%, *100nM paclitaxel+ EMD534085*: 16%, *22μM paclitaxel:* 65.22%, *22μM paclitaxel+EMD534085*: 9.5%)(**Fig. 4b,c**). Similarly, the number of dfMT-harbouring keratinocytes per animal was reduced with EMD534085/paclitaxel co-administration (*96hr vehicle*: 0±0 vs. *EMD534085*: 0±0 vs. *100nM paclitaxel*: 0.4±0.21 vs. *100nM paclitaxel+EMD534085*: 0.8±0.38 vs. *22μM paclitaxel*: 6.957±1.49 vs. 22μM *paclitaxel+EMD534085*: 0.52±0.36 cells/animal)(**Fig. 4d**). Since we administered 25μM EMD534085 after 2d of vehicle and paclitaxel treatment, the few keratinocytes harbouring dfMTs in the presence of 22μM paclitaxel and EMD534085 likely formed prior to inhibitor administration, consistent with our finding that dfMTs are already present in some animals after 48hr paclitaxel treatment (**Fig. 1e,f**).

**Figure 4.**
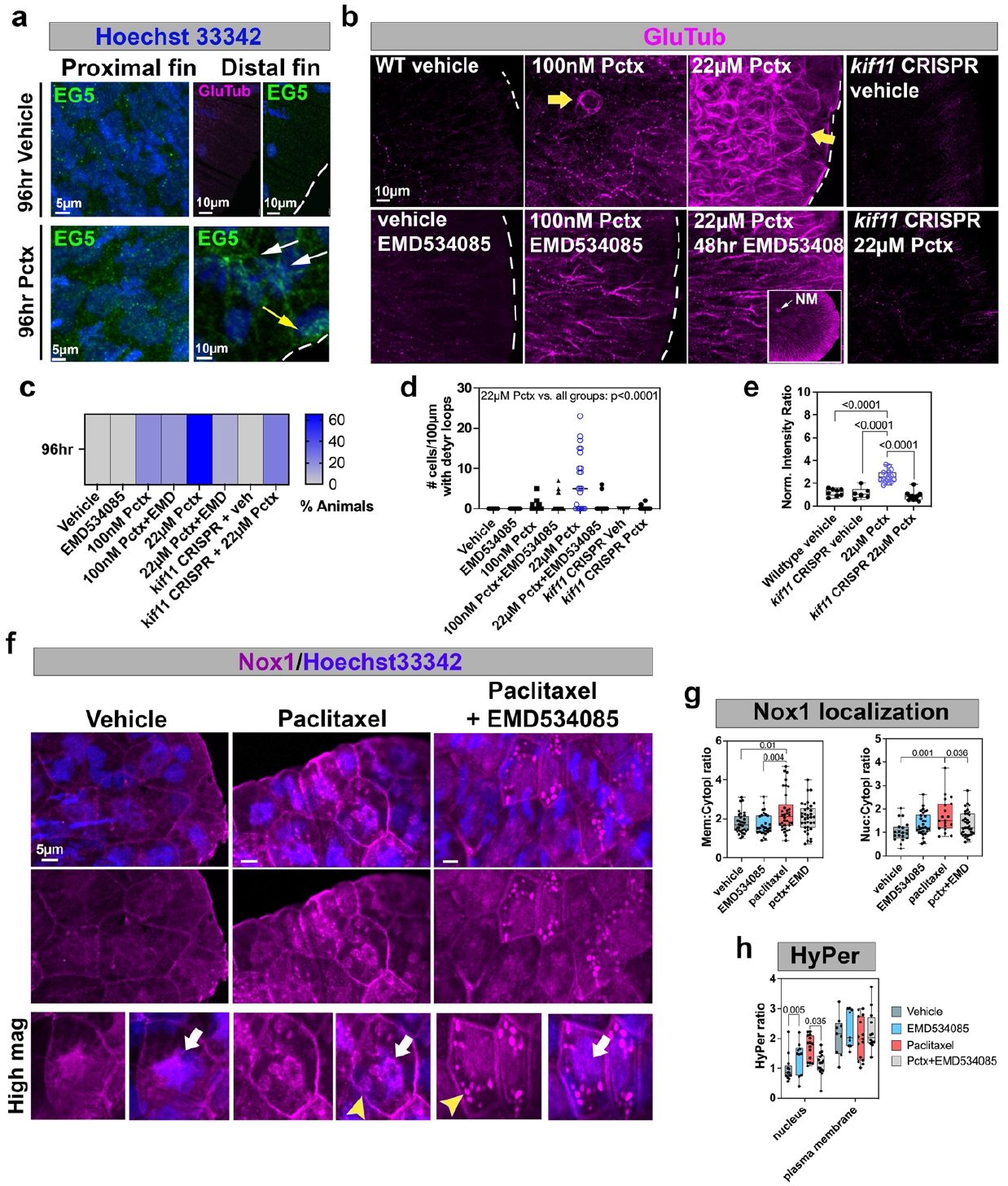
Paclitaxel induces Eg5 expression and Eg5-dependent dfMT and nuclear X-ROS formation. (**a**) Eg5 immunofluorescence staining shows punctate staining in proximal caudal fin keratinocytes of vehicle treated animals. Staining along the fin edge (white dashed line) is absent. Paclitaxel treatment promotes Eg5 activation in the proximal fin (white arrows) and in fin edge keratinocytes leading to aster-like formations (white dashed line, yellow arrow). (**b**) EMD534085 co-administration with 100nM and 22μM paclitaxel rescues microtubule fasciculation (yellow arrows) but not detyrosination. *kif11*CRISPR knockout prevents detyrosination and fasciculation of MTs. (**c**) Percentage of animals with dfMTs/keratinocyte is reduced when paclitaxel is administered in combination with EMD534085 or in *kif11* CRISPR knockout fish (n≥23 animals/group). (**d**) EMD534085 treatment or *kif11* CRISPR knockout in presence of paclitaxel reduces the number of keratinocytes with dfMTs per animal compared with paclitaxel alone (n≥21 animals per group). (**e**) Normalized intensity ratios of GluTub suggest that *kif11* CRISPR knockout rescues microtubule detyrosination induced by paclitaxel. (**f, g**) Nox1 nuclear (white arrows) but not plasma membrane translocation induced by 96hr paclitaxel treatment is rescued with EMD534085 co-administration. Vesicular Nox1 accumulation (yellow arrowheads) is prominent following paclitaxel and EMD534085 treatment. (**h**) Nuclear HyPer oxidation induced by paclitaxel treatment is rescued with EMD534085 co-administration. *Abbreviations: NM=lateral line neuromast*.

To validate the specificity of Eg5, we further transiently deleted *kif11* by injecting CRISPR oligos into 1-cell stage embryos. These were directed to exon 5 (∼90aa of 1072aa), which eliminated Eg5 protein expression (**Fig. S5**)(percent animals with dfMTs: *CRISPR*+*vehicle*: 0%; *CRISPR+22μM paclitaxel*: 25%; number of keratinocytes per animal with dfMTs: *CRISPR+vehicle*: 0±0; *CRISPR+22μM paclitaxel*: 0.29±0.16)(**Fig 4b-d**). Interestingly, transient *kif11* CRISPR knockout also reduced microtubule detyrosination when comparing 96hr vehicle and paclitaxel treated animals, suggesting a role for Eg5 in detyrosination, which was not modified when animals were treated with EMD534085 (normalized detyrosination fluorescence intensity: *vehicle:* 1.2±0.11 vs. *CRISPR+vehicle:* 1.12±0.2; *22μM paclitaxel:* 2.62±0.14 vs. *CRISPR+paclitaxel:* 0.85±0.11)(**Fig. 4e**). Importantly, increased total and phospho-EG5 levels were also detected in basal and suprabasal epidermal keratinocytes of mice following a single injection of either vehicle or 20mg/kg paclitaxel (**Fig. S6)**. These findings establish a conserved role for paclitaxel in epidermal Eg5 induction.

To determine whether Eg5 acts upstream of X-ROS and sensory axon degeneration, we first analysed the subcellular localization of Nox1 in the presence and absence of EMD534085. Immunofluorescence staining showed that EMD534085 co-administration with paclitaxel led to more condensed cytoplasmic vesicles compared with paclitaxel alone. Although EMD534085 did not affect paclitaxel-induced Nox1 plasma membrane translocation (*Membrane:cytoplasmic ratio*: vehicle: 1.78±0.09 vs. EMD534085: 1.7±0.1 vs. paclitaxel: 2.35±0.17 vs. paclitaxel+ EMD534085: 2.072±0.12), Nox1 was however largely absent from the nucleus in EMD534085-treated animals (*Nucleus:cytoplasmic ratio*: vehicle: 1.04±0.09 vs. EMD534085: 1.36±0.08 vs. paclitaxel: 1.73±0.1.6 vs. paclitaxel+ EMD534085: 1.35±0.08)(**Fig. 4f,g**). This indicates that Eg5 induces Nox1 nuclear, but not plasma membrane, translocation in a paclitaxel-dependent manner.

Further quantification of compartmentalized HyPer oxidation near the plasma membrane and the nucleus (co-labelled with Hoechst33342) showed significantly increased oxidation within the nucleus upon paclitaxel treatment, but surprisingly not at the plasma membrane. Nuclear HyPer oxidation was reduced upon co-administration of EMD534085 without significant effects on plasma membrane oxidation (**Fig. 4h**). These findings support a role for Eg5 in nuclear X-ROS production as a result of microtubule fasciculation and increased mechanical tension.

### Detyrosinated fasciculated microtubules associate with nuclei

We expected that dfMTs should be in close proximity to the nucleus in order to activate Nox1 by mechanotransduction. To further explore this possibility, we used high resolution imaging and 3D rendering of keratinocyte dfMTs and nuclei. This demonstrated that 96hr paclitaxel treatment led to a partial association between dfMTs and nuclei (**Fig. 5a, Movie S6**) whereby the nuclear content was sometimes pinched off or nuclei were perforated by dfMTs (**Fig. 5b, Movies 7,8**). Moreover, dfMTs were typically associated with significantly enlarged nuclei (vehicle nuclear volume: 291.3±30.62μm^3^ vs. paclitaxel: 582±83.18μm^3^)(**Fig. 5c**) that were less spherical (1=spherical vs. 0= nonspherical: vehicle: 0.27±0.01 vs. paclitaxel: 0.23±0.01)(**Fig. 5d**). This suggests that dfMTs form a physical barrier that impairs nuclear function.

**Figure 5.**
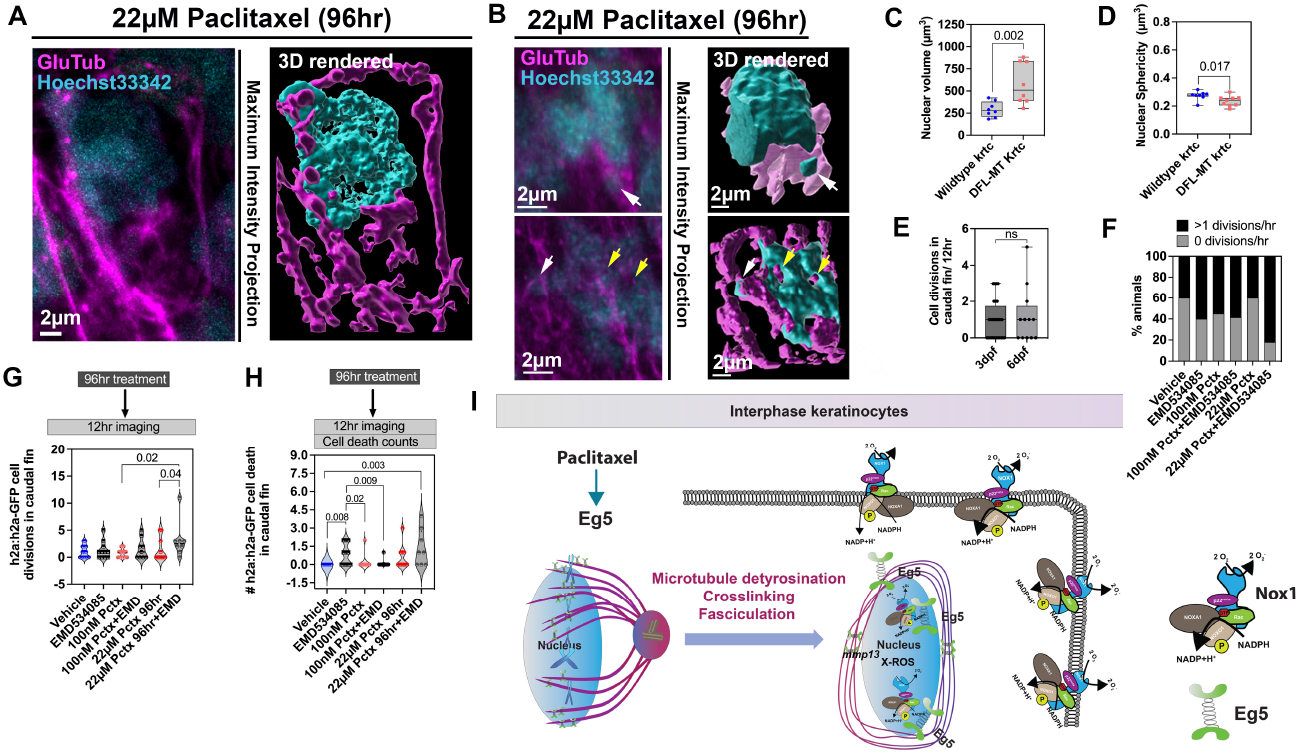
dfMTs modulate mitosis but not apoptosis. (**a**) dfMTs detected with anti-GluTub staining associate with a keratinocyte nucleus labelled with Hoechst33342 (left image). 3D reconstruction of the same image using Imaris (right image). (**b**) 3D rendered dfMTs pinch off nuclear content (left and right, white arrow) and perforate the nucleus (right, yellow arrows)(**Movies S7,8**). (**c**) Increased nuclear volume in keratinocytes harbouring dfMTs following 96hr paclitaxel (22μM) treatment compared with nuclear volume from paclitaxel-exposed cells without dfMTs (n≥5 animals/group). (**d**) Decreased nuclear sphericity in keratinocytes with dfMTs (n≥5 animals/group). (**e**) Keratinocyte divisions at 3 and 6dpf (96hr treatment) assessed by time-lapse imaging in caudal fins of Tg(*h2a*:h2a-GFP) fish (n=28 at 3dpf; n=12 at 6dpf). (**f**) Percent animals without and ≥1 mitotic divisions/12hr (n≥9 animals/group). (**g**) EMD534085 combination with 22μM paclitaxel significantly increases cell divisions/animal compared with paclitaxel treatment (n≥8 animals/group). (**h**) 96hr treatment with EMD534085 and 22μM paclitaxel significantly increases cell death (n≥8 animals/group). (**i**) Model for Eg5-dependent dfMT formation and downstream events: Paclitaxel activates Eg5 as part of the cell cycle checkpoint, however, independent of mitosis in interphase keratinocytes leading to crosslinking and fasciculation of long-term stabilized, detyrosinated microtubules. dfMTs constrain the nucleus and promote Nox1 nuclear accumulation and X-ROS formation upstream of *mmp13* induction.

### EMD534085 promotes keratinocyte mitosis and cell death in the presence of paclitaxel

Given that paclitaxel induces Eg5 expression in keratinocytes, and in cancer cells has a major role in preventing mitosis while promoting apoptosis^58-60^, we wanted to investigate whether Eg5 inhibition in combination with paclitaxel influences keratinocyte mitosis and apoptosis. If dfMTs form a physical barrier, we should see increased mitosis when EMD534085 is co-administered with paclitaxel. Consistent with this hypothesis, *in vitro* studies using paclitaxel sensitive and resistant ovarian cancer cells showed that Eg5 inhibition with HR22C16-A1 antagonized the effects of paclitaxel on mitotic inhibition^61^. To examine the effects of EMD534085 on mitosis, we quantified cell divisions in keratinocytes expressing nuclear *h2a-h2a:*GFP using 12hr time-lapse imaging. To our surprise, we found that cell divisions in caudal fin keratinocytes at 3dpf (embryonic to larval transition) and 6dpf (late larval stage) were rare. The few divisions present were mostly restricted to the distal notochord region, with an average of 0.96±0.20/12hr divisions at 3dpf and 1.16±0.44/12hr at 6dpf in this region (**Fig. 5e, Movie S9**). We found that at least 1 cell division/12hr was present in 40-60% of animals either treated with vehicle, 100nM and 22μM paclitaxel, as well as upon treatment with EMD534085 and 100nM paclitaxel/EMD534085. Combination treatment with 22μM paclitaxel and EMD534085 increased the percentage of animals with at least 1 cell division to 88% (**Fig. 5f**). Mitotic divisions per animal were also significantly increased with EMD534085 and 22μM paclitaxel in combination (96hr, *vehicle*: 1.07±0.3 vs. *100nM paclitaxel*: 0.72±0.423 vs. *22μM paclitaxel:* 1.4±0.67 vs. *EMD534085*: 1.27±0.46/12hr vs. *100nM paclitaxel+EM534085*: 1.55±0.62/12hr vs. *22μM paclitaxel+EMD534085*: 3.125±1.18)(**Fig. 5g**). This suggests that Eg5 dependent dfMT formation promotes nuclear dysfunction in keratinocytes.

Since apoptosis has been found to be the primary mechanism of cell death upon paclitaxel treatment^59-63^, we further explored the effects of 96hr paclitaxel/EMD534085 combination treatment on cell death by imaging *h2a:h2a*-GFP fish with *in vivo* time-lapse imaging for 12 hours. First, we noticed that overall, there was either none or very little cell death in vehicle and 100nM paclitaxel-treated animals, whereas 22μM paclitaxel slightly but significantly increased the number of dying cells (*vehicle*: 0±0 vs. *100nM paclitaxel*: 0.18±0.18 vs. *22μM paclitaxel*: 0.6±0.3 dying cells/ caudal fin). EMD534085 alone significantly increased cell death slightly above 22μM paclitaxel treatment, and surprisingly reduced cell death when co-administered with 100nM paclitaxel (*EMD534085*: 0.81±0.26 vs. *100nM paclitaxel+EMD534085*: 0.07±0.07). Nevertheless, EMD534085 combination with 22μM paclitaxel most significantly increased cell death compared with vehicle (*22μM paclitaxel+EMD534085*: 1.37±0.53)(**Fig. 5h,S7**). Thus, although EMD534085 in combination with a paclitaxel slightly enhances mitosis, its effects on cell death are also enhanced. These findings indicate that keratinocyte dfMTs are formed independently of mitosis but impact mitotic behaviour, while promoting apoptosis.

### EG5 does not influence microtubule growth behaviour

Our findings suggest that Eg5 acts on stable microtubules. To validate this, we characterized microtubule growth dynamics in basal keratinocytes expressing the microtubule plus-end binding protein, EB3-GFP (courtesy of R. Koester)^64^. Time-lapse recordings (1fr/sec for 60sec) showed that, as expected, EB3-GFP mostly localized to microtubule plus-ends during growth. However, a small proportion of keratinocytes also harboured uniformly labelled EB3-positive microtubules, consistent with previous observations in murine axons^65^. Paclitaxel treatment for 3hr resulted in an increased number of keratinocytes with uniform EB3 labelling (*3hr vehicle*: 5.12% vs. *paclitaxel*: 19.29%; *48hr vehicl*e: 0% vs. *paclitaxel*: 10.09%; *96hr vehicle*: 3.44% vs. 22μM *paclitaxel*: 20.13%)(**Fig. S8a,b**), indicative of microtubule stabilization. Intriguingly however, microtubules with uniform EB3-GFP binding appeared linear and continuous treatment for 48hr and 96hr with paclitaxel further enhanced the linearity (**Fig. S8c, Movie S10,11**). This behaviour therefore contrasts the curved conformation of stable dfMTs. Further analysis of microtubule growth dynamics showed that 3hr paclitaxel treatment decreased the growth velocity and track length of microtubules, whereas comet duration per se was not affected (**Fig. S8d-g**). Combination treatment of EMD534085 and 22μM paclitaxel did not significantly alter the effects induced by paclitaxel alone, suggesting that Eg5 does not influence paclitaxel-dependent microtubule growth dynamics.

To further determine whether microtubule stabilization per se leads to curved microtubules, or whether this is a specific property pertaining to paclitaxel, we analysed microtubules in keratinocytes of zebrafish injected with *CMV*:Tau-Bfp2. This microtubule-associated protein has been shown to induce microtubule stabilization by binding to the interface between α and β-tubulin heterodimers^66,67^. We found that Tau-Bfp2 induced fasciculation and mild curving of microtubules (**Fig. S9a**), which differed from the curved dfMT conformation induced by paclitaxel. Intriguingly, treatment of zebrafish expressing Tau-Bfp2 in keratinocytes for 96hr with paclitaxel promoted linearization of microtubules rather than circularization (1=straight, 0=curved: *96hr weak Tau-Bfp2 expression*: 0.94±0.01, *strong Tau-Bfp2 labelling*: 0.93±0.01, *strong Tau-Bfp2+paclitaxel*: 0.98±0.002) (**Fig. S9b,c**), consistent with previous findings^66^. Tau therefore seems to prevent paclitaxel-induced dfMT formation. Together this data suggests that dfMTs form as a consequence of paclitaxel dependent Eg5 activation rather than microtubule stabilization per se.

Interestingly, co-expression of EB3-GFP with Tau-Bfp2 in keratinocytes revealed a subpopulation of growing EB3-GFP plus-ends that grew along stable Tau-Bfp2 microtubules (**Fig. S9d, Movies S12-14**). Treatment with 22μM paclitaxel for 96hr further increased Tau-Bfp2 and EB3-GFP co-localization regardless of weak or strong Tau-Bfp2 expression (EB3-GFP puncta co-localized with Tau-Bfp2, *weak Tau-Bfp2 expression*: 85.42±1.64% of, *strong Tau-Bfp2 expression*: 86.13±1.22%, *Tau-Bfp2+*96hr *paclitaxel:* 91.26±1.64%). Also the velocity of EB3-GFP plus-ends moving along these linear Tau-Bfp2 microtubules was significantly increased with paclitaxel treatment compared to Tau-Bfp2 alone (*weak Tau-Bfp2*: 0.155±0.003μm/s, *high Tau-Bfp2*: 0.156±0.003μm/s vs. *Tau-Bfp2+96hr paclitaxel*: 0.184±0.003μm/s)(**Fig. S9e**). Plotting of the X/Y positions of some microtubules revealed that EB3-GFP growth trajectories closely followed Tau-Bfp2 associated microtubule filaments (**Fig. S9f**). Thus EB3-GFP plus-ends appear in part to grow along pre-existing stable microtubules, which could be a means to maintain cellular organization. In conclusion, the curved dfMT conformation appears to be a specific feature of paclitaxel treatment.

### Keratinocyte-specific Eg5 overexpression promotes cutaneous sensory axon degeneration

We previously reported that the earliest signs of paclitaxel-induced axon degeneration are evident at the distal fin edge due to MMP-13 dependent ECM degradation^16,17^, consistent with the vast majority of dfMTs being present in this region (**Fig. S1d**). We therefore speculated that Eg5-dependent dfMTs induce axon degeneration. To test this, we first analysed axon degeneration in zebrafish that were treated for 96hr with low (100nM) and high (22μM) paclitaxel concentrations in the presence and absence of EMD534085. Treatment with vehicle, EMD534085 and 100nM paclitaxel did not alter axon branch number assessed with acetylated tubulin staining. Treatment with 22μM paclitaxel, however, significantly reduced the axon branch number as previously shown (*96hr: vehicle*: 10.83±0.58, *EMD534085:* 10.25±0.47, *100nM paclitaxel*: 9.86±0.43,*100nM paclitaxel+48hr EMD534085:* 10.73±0.63, *22μM paclitaxel:* 7.54±0.56 branches/50μm)(**Fig. 6a,b**). Intriguingly, combination treatment of 22μM paclitaxel and EMD534085 significantly rescued axon degeneration even above control levels (19.89±1.09 branches/50μm). These findings are consistent with observations that monastrol promotes neurite outgrowth in cultured mouse DRG neurons^68^, suggesting a neuron-intrinsic growth effect. To further validate Eg5 as the pharmacological target of EMD534085 in axon regeneration, we also analysed zebrafish injected with *kif11* CRISPR oligo. Paclitaxel-treated *kif11* CRISPR zebrafish did not show altered axon branch numbers in the caudal fin compared with vehicle-treated CRISPR and wildtype controls, although axon growth was not increased, as seen with EMD534085 (*kif11-CRISPR+vehicle*: 12.08±0.68, *kif11-CRISPR+22μM paclitaxel*: 12.83±0.96)(**Fig. 6b**). These findings implicate Eg5 as downstream target of paclitaxel in axon degeneration. It remains to be investigated why *kif11* knockout modulates axon behaviour differently from pharmacological Eg5 inhibition, possibly due to developmental effects of the CRISPR oligo.

**Fig. 6.**
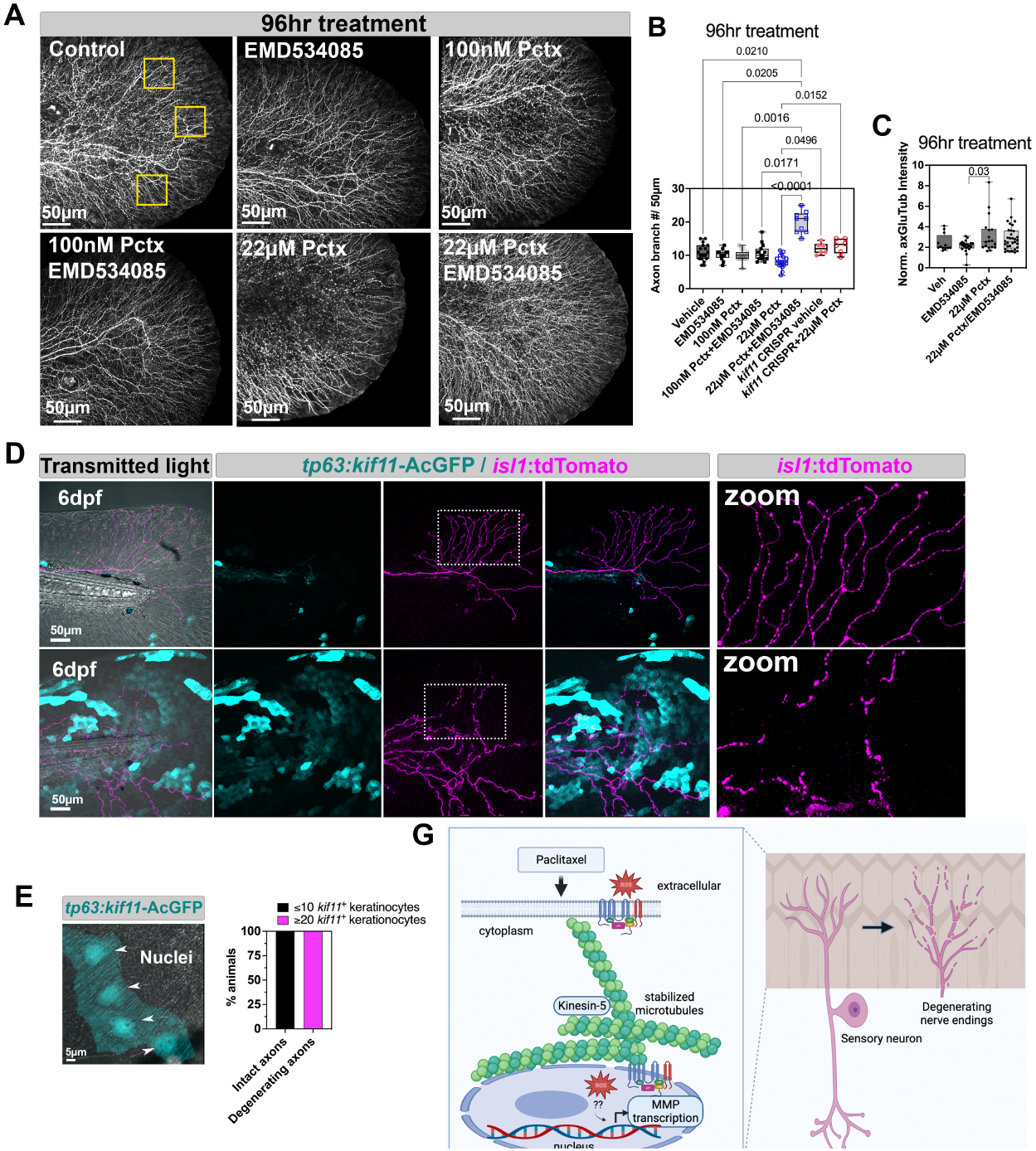
Eg5 inhibition rescues and keratinocyte-specific overexpression induces paclitaxel neurotoxicity. (**A**) Degeneration of cutaneous axons detected with anti-acetylated tubulin antibody staining following 96hr treatment with 22μM paclitaxel, but not when treated with 100nM paclitaxel or 22μM paclitaxel+EMD534085 (n≥11 animals/group). (**B**) Quantification shows an increased axon branch number for paclitaxel+EMD534085, whereas *kif11* CRISPR knockout rescues axon branch number to wildtype levels (n=6 animals/group). (**C**) Significantly increased axonal dMTs along axon segments in distal caudal fin seen with 22μM paclitaxel compared to EMD534085 (n=5-7 animals/group). (**D**) Top panel: Co-expression of *isl1:*LexA-lexaop_14xUAS-tdTomato (magenta) in axons and *tp63*:*kif11*-AcGFP (blue) in few keratinocytes that are not in contact with cutaneous branches does not promote axon degeneration. Bottom panel: Axon degeneration is prominent when *tp63*:*kif11*-AcGFP expressing basal keratinocytes are abundant and establish contact with cutaneous branches. (**E**) Magnification of (d) shows nuclear Eg5 and weaker cytoplasmic localization in 6dpf zebrafish expressing *tp63*:*kif11*-AcGFP. (**F**) Quantification of axon degeneration shows low presence of keratinocytes expressing *kif11*-AcGFP is insufficient to induce axon degeneration. (**G**) Model showing paclitaxel-dependent Eg5 induction leading to dfMT formation and Nox1-dependent *mmp13* expression in keratinocytes, which promotes ECM degradation and axon degeneration.

We next wanted to determine if Eg5-dependent axon degeneration was caused by keratinocyte or neuron-specific effects given that Eg5 is also expressed neuronally^69,70^ and we observed possible neuron-intrinsic axon growth effects with EMD534085. We initially determined to what extent axonal microtubules are stabilized and whether this process is altered using EMD534085 in the presence and absence of paclitaxel.

Quantification of axonal detyrosination in wildtype distal fins following 96hr treatment with either vehicle or EMD534085 showed a punctate pattern in axons, whereas 22μM paclitaxel but not paclitaxel/EMD534085 promoted a uniform distribution of dMTs along axon segments (**Fig. S10a**). Quantification of the average detyrosination fluorescence intensity along individual axon segments however showed that only paclitaxel significantly differed from EMD534085 (normalized GluTub intensity ratios, wildtype 96hr *vehicle*: 2.44±0.24, *EMD534085:* 2.13±0.13, *22μM paclitaxel*: 3.19±0.42, *22μM paclitaxel+EMD534085:* 2.86±0.21)(**Fig. 6c, S10a**). Thus, paclitaxel does not appear to regulate dMT formation via Eg5 in axons.

We next injected a transgene encoding *tp63*:*kif11*-AcGFP for transient Eg5 overexpression in keratinocytes. We further co-expressed *isl1*:tdTomato (gift from Alvaro Sagasti, UCLA) in sensory neurons to label wildtype axons. Eg5 preferentially located to keratinocyte nuclei with some cytoplasmic labelling (**Fig. 6d,e**), consistent with its nuclear localization in cultured interphase HeLa cells^71^. Transient plasmid DNA injections into zebrafish typically mosaically label cells, which allowed for the observation of axons in the absence and presence of Eg5-overexpressing keratinocytes. Whereas wildtype axons in animals with few Eg5-GFP positive (≤10) keratinocytes) remained intact (**Fig. 6d**, top panel), axons in contact with large areas of Eg5 overexpressing keratinocytes (≥20) degenerated (**Fig. 6d** (bottom panel), **Fig. 6f**). Therefore, keratinocyte-specific Eg5 overexpression mimics paclitaxel-induced sensory axon degeneration. These findings support a model in which Eg5 activation by paclitaxel stimulates X-ROS formation and MMP-13 dependent ECM degradation in keratinocytes, ultimately leading to axon degeneration (**Fig. 6g**).

## Discussion

### Paclitaxel treatment induces ectopic Eg5 expression in non-mitotic cells

We have shown that paclitaxel ectopically activates Eg5 and cell cycle regulatory genes in largely non-mitotic keratinocytes, suggesting that paclitaxel triggers the expression or activity of these genes regardless of cell cycle status and thus explains its side effects. Given the role of Eg5 in mitosis, ectopically activated Eg5 appears to promote microtubule crosslinking but due to the lack of mitotic signals, mitosis does not proceed and instead leads to dfMT formation. How Eg5 is activated by paclitaxel remains unclear. In yeast, the cell cycle regulator, Mad1 recruits Cut7 (Eg5) to kinetochores of misaligned chromosomes and promotes chromosome gliding towards the spindle equator^72^. A parallel mechanism triggered by paclitaxel treatment may therefore activate Eg5 binding to microtubules, which remains to be tested. It is also possible that post-translational modifications, such as the acetylation of K146 in Eg5 leads to its activation. Studies have shown that K146 serves as on/off switch for Eg5 motor activity^71^. Our own studies show that paclitaxel treatment increases keratinocyte-specific acetylation of alpha-tubulin (K40) at the distal fin edge where detyrosination is evident (**Fig. S10**), consistent with this possibility.

### Eg5 regulates Nox1 activity via mechanotransduction mechanisms

We discovered that paclitaxel promotes Nox1 translocation to the plasma membrane and nucleus, but that Eg5 only influences nuclear X-ROS formation whereas no increase in H_2_O_2_ was detected at the plasma membrane. One possibility is that since O_2_^-^ and H_2_O_2_ are secreted into the extracellular space by NADPH oxidases, mechanisms that promote diffusion into keratinocytes could be absent, or Nox1 translocation occurs independent of its activation given that it requires various subunits for activation^44,73^. Since HyPer is selective for H_2_O_2_, there may also be other ROS formed at the plasma membrane, which we did not capture. Nuclear H_2_O_2_ formation, however, is consistent with the H_2_O_2_-dependent activation of MMPs^74^, and supports a model by which nuclear X-ROS may locally induce *MMP13* transcription. Although the molecular mechanisms underlying these observations have not been established, known MMP-13 transcriptional regulators, such as AP-1 and MAP kinases may mediate this process^75,76^. Proteins that link microtubules to the nucleus might also be activated. For instance, Rac1 is a critical regulator of both cytoskeletal dynamics and Nox2 activity^77^. Rac1 can bind IQGAP1, a protein that tethers actin filaments and microtubules to the cell periphery at the plasma membrane and to the face of the nuclear envelop^78,79^, consistent with paclitaxel dependent Nox1 translocation to the plasma membrane and nucleus in keratinocytes. It also remains unclear how Eg5 inhibition promotes Nox1 nuclear exclusion and accumulation in cytoplasmic vesicles. Consistent with our findings, Nox enzymes have been detected in exosomes, such as during transport from macrophages into dorsal root ganglion neurons^80^, and in extracellular vesicles, whereby Nox release was associated with inflammation in platelets^81^. Given that NADPH oxidases are membrane-integral enzyme complexes, vesicular transport appears to promote their intra- and extracellular motility, for which microtubules may be the primary regulators in an Eg5 dependent manner under conditions of paclitaxel treatment.

### Increased microtubule detyrosination is correlated with disease

Detyrosinated microtubules provide mechanical resistance to cells, such as during cardiac contraction, but they have been primarily associated with pathological conditions. For instance, patients with heart failure display an increased load of detyrosinated microtubules in cardiomyocytes^23^, which slows the contraction–relaxation cycle normally found in healthy cardiomyocytes^22^. Our findings support the idea that microtubule detyrosination is associated with pathological changes in cells. Detyrosination pathology, however, seems to be cell type-dependent since axons showed an increase in microtubule detyrosination when paclitaxel+EMD534085 were co-administered despite that axon degeneration was prevented and axon growth promoted. Nevertheless, EMD534085 promoted axon growth in the presence of paclitaxel unlike *kif11* knockout, which rescued paclitaxel-induced axon degeneration but did not promote growth. It therefore appears that short-term and chronic microtubule manipulations by Eg5 in axons have distinct effects and are not involved in axonal pathology, in contrast to keratinocytes.

### ZStretcher dependent ROS production

We have shown that HyPer oxidation differs between stretched vehicle control and paclitaxel treated wildtype animals, whereas no notable difference was observed for *cyba* mutants. In these experiments, paclitaxel treatment was performed for only 3hr since we observed changes in microtubule growth dynamics even after a short treatment. However, longer treatment periods may cause different effects, which remains to be investigated. Nevertheless, our results conform to the hypothesis that paclitaxel-dependent microtubule stabilisation alters ROS production, consistent with studies in muscle cells^23,24,31-34^. We previously demonstrated that axons do not display increased ROS levels^17^, and since ROS diffusion from adjacent cell types is relatively short-lived, it seems unlikely that measured HyPer oxidation levels in keratinocytes are related to ROS production in other cell types. Although other cells and ECM molecules are also stretched, the lack of oxidation at the plasma membrane suggests that H2O2 doesn’t easily diffuse into cells if it was produced in the vicinity of keratinocytes. Thus, the differences in HyPer oxidation in the presence of paclitaxel point to a role for microtubule mediated Nox activation.

### Tau and EB3 overexpression in keratinocytes promote microtubule linearization

We showed that Tau and EB3 overexpression in keratinocytes promoted the linearization of microtubules regardless of paclitaxel presence, consistent with *in vitro* observations^66,82^. Linearization of microtubules may be a direct consequence of cooperative binding between paclitaxel and Tau, whereby paclitaxel binding sites are occupied in the presence of Tau^67^. The looped conformation of dfMTs present in keratinocytes following paclitaxel treatment may be a specific consequence of Eg5 activation, whereas Eg5 may not be induced by paclitaxel in the presence of Tau or its binding sites on microtubules are occupied by Tau.

### Cell crowding may contribute to paclitaxel neurotoxicity

The appearance of dfMTs primarily in fin edge keratinocytes closely correlates with the onset of paclitaxel neurotoxicity in this region following prolonged paclitaxel treatment^16^. The distal fin edge is under high tension, which promotes the extrusion of live and apoptotic cells under crowding conditions^83^. Our previous studies suggested that the fin edge may undergo less remodelling in the presence of paclitaxel due to MMP-13 dependent ECM degradation that promotes epidermal abrasions^17^. One hypothesis to be tested is that fin edge keratinocytes are no longer extruded in the presence of paclitaxel due to the altered mechanical properties of keratinocytes, which induces axon degeneration. A parallel process could take place in mammalian skin where the lack of differentiating keratinocytes prevents their movement toward the surface, leading to axon damage.

## Supporting information

Mesenchymal cell morphology

Detyrosinated microtubules (anti-GluTub) and nuclei (Hoechst 33342) in the caudal fin of a 96hr vehicle-treated zebrafish (6dpf

Detyrosinated microtubules detected with anti-GluTub antibody staining and nuclei detected with Hoechst 33342 staining in the caudal fin of

3D reconstruction of Nox1 staining in a single epidermal keratinocyte following vehicle treatment

3D reconstruction of Nox1 staining in a single epidermal keratinocyte following paclitaxel treatment

3D reconstruction of a nucleus and surrounding dfMTs in a caudal fin keratinocyte following paclitaxel treatment

3D reconstruction of dfMTs pinching off nuclear content

Fluorescence staining and 3D reconstruction of a nucleus that is perforated by dfMTs

Live imaging of cell division

Plus-end tracking of keratinocyte microtubules in a vehicle-treated zebrafish

EB3 is uniformly distributed along microtubules following paclitaxel treatment

Co-expression of tp63:EB3-GFP and CMV:Tau-Bfp2 in a vehicle-treated fish

Co-expression of tp63:EB3-GFP and CMV:Tau-Bfp2 in a paclitaxel-treated fish

3D reconstruction of EB3-GFP and Tau-Bfp2 following paclitaxel treatment

## Author contributions

A.C.D. and T.S.L. performed experiments and edited the manuscript.

M.X.X. supervised the mouse studies and edited the manuscript.

A.M.C., A.D.P., N.A.S., M.J.U., L.R.K.R., L.G.P. and M.C.A.S. performed experiments and analysed data.

D.P.R. and C.H. engineered the ZStretcher and wrote the code for the software-controlled motor unit.

S.H. and N.P.S. conducted the human studies.

S.W. wrote the python code and generated the polar plots.

S.R. directed the studies, performed experiments, analysed the data, and wrote and edited the manuscript.

## Acknowledgements

We thank Ricardo Cepeda and Dr. Julia Dallman for zebrafish maintenance, and Drs. Alvaro Sagasti and Reinhard Koester for providing reagents. We also thank Dr. James Baker for assisting with imaging. We further thank the OpenSource software developers of ImageJ/ Fiji for their contributions with various plugins, and Bitplane, Micro Optics of Florida (Rob Celestine and Ron Smircich), and Zeiss (Oliver Tress) for their generous support, as well as Addgene donors for their plasmid contributions. This research was supported by funds from NIH R01CA215973-05, P20GM103423 NIH-NIGMS NCATS CTSI Miami, NIH-NCI, R21 NS094939 NIH-NINDS, P20GM104318 NIH-NIGMS.

## Competing interests

The authors have no competing interests in the research.

## Notes

### Competing Interest Statement

The authors have declared no competing interest.

### Summary of Updates

Change of competing interest statement declaring no competing interests.

## References

1 Gornstein, E. & Schwarz, T. L. The paradox of paclitaxel neurotoxicity: Mechanisms and unanswered questions. Neuropharmacology 76 Pt A, 175–183, doi:10.1016/j.neuropharm.2013.08.016 (2014).

2 Argyriou, A. A., Bruna, J., Genazzani, A. A. & Cavaletti, G. Chemotherapy-induced peripheral neurotoxicity: management informed by pharmacogenetics. Nat Rev Neurol 13, 492–504, doi:10.1038/nrneurol.2017.88 (2017).

3 Argyriou, A. A., Bruna, J., Marmiroli, P. & Cavaletti, G. Chemotherapy-induced peripheral neurotoxicity (CIPN): an update. Crit Rev Oncol Hematol 82, 51–77, doi:10.1016/j.critrevonc.2011.04.012 (2012).

4 Argyriou, A. A. et al. Neurophysiological, nerve imaging and other techniques to assess chemotherapy-induced peripheral neurotoxicity in the clinical and research settings. Journal of neurology, neurosurgery, and psychiatry, doi:10.1136/jnnp-2019-320969 (2019).

5 Argyriou, A. A., Zolota, V., Kyriakopoulou, O. & Kalofonos, H. P. Toxic peripheral neuropathy associated with commonly used chemotherapeutic agents. J BUON 15, 435–446 (2010).

6 Beijers, A. J., Jongen, J. L. & Vreugdenhil, G. Chemotherapy-induced neurotoxicity: the value of neuroprotective strategies. Neth J Med 70, 18–25 (2012).

7 Boyette-Davis, J. A., Walters, E. T. & Dougherty, P. M. Mechanisms involved in the development of chemotherapy-induced neuropathy. Pain Manag 5, 285–296, doi:10.2217/pmt.15.19 (2015).

8 Flatters, S. J. L., Dougherty, P. M. & Colvin, L. A. Clinical and preclinical perspectives on Chemotherapy-Induced Peripheral Neuropathy (CIPN): a narrative review. Br J Anaesth 119, 737–749, doi:10.1093/bja/aex229 (2017).

9 Bennett, G. J., Liu, G. K., Xiao, W. H., Jin, H. W. & Siau, C. Terminal arbor degeneration--a novel lesion produced by the antineoplastic agent paclitaxel. Eur J Neurosci 33, 1667–1676, doi:10.1111/j.1460-9568.2011.07652.x (2011).

10 Staff, N. P. et al. Pathogenesis of paclitaxel-induced peripheral neuropathy: A current review of in vitro and in vivo findings using rodent and human model systems. Experimental neurology 324, 113121, doi:10.1016/j.expneurol.2019.113121 (2020).

11 Seretny, M. et al. Incidence, prevalence, and predictors of chemotherapy-induced peripheral neuropathy: A systematic review and meta-analysis. Pain 155, 2461–2470, doi:10.1016/j.pain.2014.09.020 (2014).

12 Pease-Raissi, S. E. et al. Paclitaxel Reduces Axonal Bclw to Initiate IP3R1-Dependent Axon Degeneration. Neuron 96, 373–386 e376, doi:10.1016/j.neuron.2017.09.034 (2017).

13 Bobylev, I. et al. Paclitaxel inhibits mRNA transport in axons. Neurobiol Dis 82, 321–331, doi:10.1016/j.nbd.2015.07.006 (2015).

14 Li, Y. et al. The Cancer Chemotherapeutic Paclitaxel Increases Human and Rodent Sensory Neuron Responses to TRPV1 by Activation of TLR4. J Neurosci 35, 13487–13500, doi:10.1523/JNEUROSCI.1956-15.2015 (2015).

15 Brandolini, L. et al. CXCR1/2 pathways in paclitaxel-induced neuropathic pain. Oncotarget 8, 23188–23201, doi:10.18632/oncotarget.15533 (2017).

16 Lisse, T. S. et al. Paclitaxel-induced epithelial damage and ectopic MMP-13 expression promotes neurotoxicity in zebrafish. Proceedings of the National Academy of Sciences of the United States of America 113, E2189–2198, doi:10.1073/pnas.1525096113 (2016).

17 Cirrincione, A. M. et al. Paclitaxel-induced peripheral neuropathy is caused by epidermal ROS and mitochondrial damage through conserved MMP-13 activation. Sci Rep 10, 3970, doi:10.1038/s41598-020-60990-8 (2020).

18 Shin, G. J. et al. Integrins protect sensory neurons in models of paclitaxel-induced peripheral sensory neuropathy. Proceedings of the National Academy of Sciences of the United States of America 118, doi:10.1073/pnas.2006050118 (2021).

19 Hyman, A. A., Salser, S., Drechsel, D. N., Unwin, N. & Mitchison, T. J. Role of GTP hydrolysis in microtubule dynamics: information from a slowly hydrolyzable analogue, GMPCPP. Mol Biol Cell 3, 1155–1167, doi:10.1091/mbc.3.10.1155 (1992).

20 Arnal, I. & Wade, R. H. How does taxol stabilize microtubules? Curr Biol 5, 900–908 (1995).

21 Hallak, M. E., Rodriguez, J. A., Barra, H. S. & Caputto, R. Release of tyrosine from tyrosinated tubulin. Some common factors that affect this process and the assembly of tubulin. FEBS letters 73, 147–150, doi:10.1016/0014-5793(77)80968-x (1977).

22 Nieuwenhuis, J. & Brummelkamp, T. R. The Tubulin Detyrosination Cycle: Function and Enzymes. Trends Cell Biol 29, 80–92, doi:10.1016/j.tcb.2018.08.003 (2019).

23 Chen, C. Y. et al. Suppression of detyrosinated microtubules improves cardiomyocyte function in human heart failure. Nature medicine 24, 1225–1233, doi:10.1038/s41591-018-0046-2 (2018).

24 Kerr, J. P. et al. Detyrosinated microtubules modulate mechanotransduction in heart and skeletal muscle. Nature communications 6, 8526, doi:10.1038/ncomms9526 (2015).

25 Mialhe, A. et al. Tubulin detyrosination is a frequent occurrence in breast cancers of poor prognosis. Cancer research 61, 5024–5027 (2001).

26 Peris, L. et al. Tubulin tyrosination regulates synaptic function and is disrupted in Alzheimer’s disease. Brain : a journal of neurology, doi:10.1093/brain/awab436 (2022).

27 Dalous, J. et al. Reversal of cell polarity and actin-myosin cytoskeleton reorganization under mechanical and chemical stimulation. Biophys J 94, 1063–1074, doi:10.1529/biophysj.107.114702 (2008).

28 Thery, M., Jimenez-Dalmaroni, A., Racine, V., Bornens, M. & Julicher, F. Experimental and theoretical study of mitotic spindle orientation. Nature 447, 493–496, doi:10.1038/nature05786 (2007).

29 Engler, A. J., Sen, S., Sweeney, H. L. & Discher, D. E. Matrix elasticity directs stem cell lineage specification. Cell 126, 677–689, doi:10.1016/j.cell.2006.06.044 (2006).

30 Grote, K. et al. Mechanical stretch enhances mRNA expression and proenzyme release of matrix metalloproteinase-2 (MMP-2) via NAD(P)H oxidase-derived reactive oxygen species. Circulation research 92, e80–86, doi:10.1161/01.RES.0000077044.60138.7C (2003).

31 Khairallah, R. J. et al. Microtubules underlie dysfunction in duchenne muscular dystrophy. Sci Signal 5, ra56, doi:10.1126/scisignal.2002829 (2012).

32 Prosser, B. L., Ward, C. W. & Lederer, W. J. X-ROS signaling: rapid mechano-chemo transduction in heart. Science 333, 1440–1445, doi:10.1126/science.1202768 (2011).

33 Robison, P. et al. Detyrosinated microtubules buckle and bear load in contracting cardiomyocytes. Science 352, aaf0659, doi:10.1126/science.aaf0659 (2016).

34 Ward, C. W., Prosser, B. L. & Lederer, W. J. Mechanical stretch-induced activation of ROS/RNS signaling in striated muscle. Antioxid Redox Signal 20, 929–936, doi:10.1089/ars.2013.5517 (2014).

35 Roll-Mecak, A. The Tubulin Code in Microtubule Dynamics and Information Encoding. Developmental cell 54, 7–20, doi:10.1016/j.devcel.2020.06.008 (2020).

36 Khawaja, S., Gundersen, G. G. & Bulinski, J. C. Enhanced stability of microtubules enriched in detyrosinated tubulin is not a direct function of detyrosination level. The Journal of cell biology 106, 141–149, doi:10.1083/jcb.106.1.141 (1988).

37 Cassereau, J., Codron, P. & Funalot, B. Inherited peripheral neuropathies due to mitochondrial disorders. Revue neurologique 170, 366–374, doi:10.1016/j.neurol.2013.11.005 (2014).

38 Flatters, S. J. The contribution of mitochondria to sensory processing and pain. Prog Mol Biol Transl Sci 131, 119–146, doi:10.1016/bs.pmbts.2014.12.004 (2015).

39 Pareyson, D., Piscosquito, G., Moroni, I., Salsano, E. & Zeviani, M. Peripheral neuropathy in mitochondrial disorders. Lancet Neurol 12, 1011–1024, doi:10.1016/S1474-4422(13)70158-3 (2013).

40 Bilan, D. S. & Belousov, V. V. In Vivo Imaging of Hydrogen Peroxide with HyPer Probes. Antioxid Redox Signal 29, 569–584, doi:10.1089/ars.2018.7540 (2018).

41 Bilan, D. S. et al. HyPer-3: a genetically encoded H(2)O(2) probe with improved performance for ratiometric and fluorescence lifetime imaging. ACS Chem Biol 8, 535–542, doi:10.1021/cb300625g (2013).

42 Cirrincione, A., Reimonn, C., Harrison, B. & Rieger, S. Longitudinal RNA sequencing of skin and DRG neurons in mice with paclitaxel-induced peripheral neuropathy. Data 7, 72, doi:https://doi.org/10.3390/data7060072 (2022).

43 Donkó, A., Péterfi, Z., Sum, A., Leto, T. & Geiszt, M. Dual oxidases. Philos Trans R Soc Lond B Biol Sci 360, 2301–2308, doi:L7520011N0244316 [pii] 10.1098/rstb.2005.1767 (2005).

44 Bedard, K. & Krause, K. H. The NOX family of ROS-generating NADPH oxidases: physiology and pathophysiology. Physiological reviews 87, 245–313 (2007).

45 Ushio-Fukai, M., Zafari, A. M., Fukui, T., Ishizaka, N. & Griendling, K. K. p22phox is a critical component of the superoxide-generating NADH/NADPH oxidase system and regulates angiotensin II-induced hypertrophy in vascular smooth muscle cells. The Journal of biological chemistry 271, 23317–23321, doi:10.1074/jbc.271.38.23317 (1996).

46 Szklarczyk, D. et al. The STRING database in 2021: customizable protein-protein networks, and functional characterization of user-uploaded gene/measurement sets. Nucleic Acids Res 49, D605–D612, doi:10.1093/nar/gkaa1074 (2021).

47 Leonard, S. E. & Carroll, K. S. Chemical ‘omics’ approaches for understanding protein cysteine oxidation in biology. Current opinion in chemical biology 15, 88–102, doi:10.1016/j.cbpa.2010.11.012 (2011).

48 Waters, J. C., Chen, R. H., Murray, A. W. & Salmon, E. D. Localization of Mad2 to kinetochores depends on microtubule attachment, not tension. The Journal of cell biology 141, 1181–1191, doi:10.1083/jcb.141.5.1181 (1998).

49 Jia, L., Li, B. & Yu, H. The Bub1-Plk1 kinase complex promotes spindle checkpoint signalling through Cdc20 phosphorylation. Nature communications 7, 10818, doi:10.1038/ncomms10818 (2016).

50 Hirokawa, N., Noda, Y., Tanaka, Y. & Niwa, S. Kinesin superfamily motor proteins and intracellular transport. Nat Rev Mol Cell Biol 10, 682–696, doi:10.1038/nrm2774 (2009).

51 Leary, A. et al. Successive Kinesin-5 Microtubule Crosslinking and Sliding Promote Fast, Irreversible Formation of a Stereotyped Bipolar Spindle. Curr Biol 29, 3825-3837.e3823, doi:10.1016/j.cub.2019.09.030 (2019).

52 Bobylev, I. et al. Kinesin-5 Blocker Monastrol Protects Against Bortezomib-Induced Peripheral Neurotoxicity. Neurotox Res 32, 555–562, doi:10.1007/s12640-017-9760-7 (2017).

53 Brandes, R. P., Weissmann, N. & Schroder, K. Nox family NADPH oxidases in mechano-transduction: mechanisms and consequences. Antioxid Redox Signal 20, 887–898, doi:10.1089/ars.2013.5414 (2014).

54 Ge, S. X., Son, E. W. & Yao, R. iDEP: an integrated web application for differential expression and pathway analysis of RNA-Seq data. BMC Bioinformatics 19, 534, doi:10.1186/s12859-018-2486-6 (2018).

55 Meaders, J. L., de Matos, S. N. & Burgess, D. R. A Pushing Mechanism for Microtubule Aster Positioning in a Large Cell Type. Cell Rep 33, 108213, doi:10.1016/j.celrep.2020.108213 (2020).

56 Goulet, A. & Moores, C. New insights into the mechanism of force generation by kinesin-5 molecular motors. Int Rev Cell Mol Biol 304, 419–466, doi:10.1016/B978-0-12-407696-9.00008-7 (2013).

57 Mann, B. J. & Wadsworth, P. Kinesin-5 Regulation and Function in Mitosis. Trends Cell Biol 29, 66–79, doi:10.1016/j.tcb.2018.08.004 (2019).

58 Jordan, M. A., Toso, R. J., Thrower, D. & Wilson, L. Mechanism of mitotic block and inhibition of cell proliferation by taxol at low concentrations. Proceedings of the National Academy of Sciences of the United States of America 90, 9552–9556 (1993).

59 Zasadil, L. M. et al. Cytotoxicity of paclitaxel in breast cancer is due to chromosome missegregation on multipolar spindles. Sci Transl Med 6, 229ra243, doi:10.1126/scitranslmed.3007965 (2014).

60 Zhao, S., Tang, Y., Wang, R. & Najafi, M. Mechanisms of cancer cell death induction by paclitaxel: an updated review. Apoptosis : an international journal on programmed cell death 27, 647–667, doi:10.1007/s10495-022-01750-z (2022).

61 Marcus, A. I. et al. Mitotic kinesin inhibitors induce mitotic arrest and cell death in Taxol-resistant and - sensitive cancer cells. The Journal of biological chemistry 280, 11569–11577, doi:10.1074/jbc.M413471200 (2005).

62 Ren, X. et al. Paclitaxel suppresses proliferation and induces apoptosis through regulation of ROS and the AKT/MAPK signaling pathway in canine mammary gland tumor cells. Mol Med Rep 17, 8289–8299, doi:10.3892/mmr.2018.8868 (2018).

63 Jordan, M. A. et al. Mitotic block induced in HeLa cells by low concentrations of paclitaxel (Taxol) results in abnormal mitotic exit and apoptotic cell death. Cancer research 56, 816–825 (1996).

64 Distel, M., Hocking, J. C., Volkmann, K. & Köster, R. W. The centrosome neither persistently leads migration nor determines the site of axonogenesis in migrating neurons in vivo. The Journal of cell biology 191, 875–890, doi:10.1083/jcb.201004154 (2010).

65 Kleele, T. et al. An assay to image neuronal microtubule dynamics in mice. Nature communications 5, 4827, doi:10.1038/ncomms5827 (2014).

66 Kadavath, H. et al. Tau stabilizes microtubules by binding at the interface between tubulin heterodimers. Proceedings of the National Academy of Sciences of the United States of America 112, 7501–7506, doi:10.1073/pnas.1504081112 (2015).

67 Ross, J. L., Santangelo, C. D., Makrides, V. & Fygenson, D. K. Tau induces cooperative Taxol binding to microtubules. Proceedings of the National Academy of Sciences of the United States of America 101, 12910–12915, doi:10.1073/pnas.0402928101 (2004).

68 Haque, S. A., Hasaka, T. P., Brooks, A. D., Lobanov, P. V. & Baas, P. W. Monastrol, a prototype anti-cancer drug that inhibits a mitotic kinesin, induces rapid bursts of axonal outgrowth from cultured postmitotic neurons. Cell Motil Cytoskeleton 58, 10–16, doi:10.1002/cm.10176 (2004).

69 Falnikar, A., Tole, S. & Baas, P. W. Kinesin-5, a mitotic microtubule-associated motor protein, modulates neuronal migration. Mol Biol Cell 22, 1561–1574, doi:10.1091/mbc.E10-11-0905 (2011).

70 Wei, N. et al. Inhibitions and Down-Regulation of Motor Protein Eg5 Expression in Primary Sensory Neurons Reveal a Novel Therapeutic Target for Pathological Pain. Neurotherapeutics 19, 1401–1413, doi:10.1007/s13311-022-01263-2 (2022).

71 Muretta, J. M. et al. A posttranslational modification of the mitotic kinesin Eg5 that enhances its mechanochemical coupling and alters its mitotic function. Proceedings of the National Academy of Sciences of the United States of America 115, E1779–E1788, doi:10.1073/pnas.1718290115 (2018).

72 Akera, T., Goto, Y., Sato, M., Yamamoto, M. & Watanabe, Y. Mad1 promotes chromosome congression by anchoring a kinesin motor to the kinetochore. Nat Cell Biol 17, 1124–1133, doi:10.1038/ncb3219 (2015).

73 Clark, R. A., Epperson, T. K. & Valente, A. J. Mechanisms of activation of NADPH oxidases. Jpn J Infect Dis 57, S22–23 (2004).

74 Cook-Mills, J. M. Hydrogen peroxide activation of endothelial cell-associated MMPs during VCAM-1-dependent leukocyte migration. Cell Mol Biol (Noisy-le-grand) 52, 8–16 (2006).

75 Son, Y. et al. Mitogen-Activated Protein Kinases and Reactive Oxygen Species: How Can ROS Activate MAPK Pathways? J Signal Transduct 2011, 792639, doi:10.1155/2011/792639 (2011).

76 Dhar, A., Young, M. R. & Colburn, N. H. The role of AP-1, NF-kappaB and ROS/NOS in skin carcinogenesis: the JB6 model is predictive. Molecular and cellular biochemistry 234-235, 185–193 (2002).

77 D’Ambrosi, N., Rossi, S., Gerbino, V. & Cozzolino, M. Rac1 at the crossroad of actin dynamics and neuroinflammation in Amyotrophic Lateral Sclerosis. Front Cell Neurosci 8, 279, doi:10.3389/fncel.2014.00279 (2014).

78 Johnson, M. A. & Henderson, B. R. The scaffolding protein IQGAP1 co-localizes with actin at the cytoplasmic face of the nuclear envelope: implications for cytoskeletal regulation. Bioarchitecture 2, 138–142, doi:10.4161/bioa.21182 (2012).

79 Owen, D. et al. The IQGAP1-Rac1 and IQGAP1-Cdc42 interactions: interfaces differ between the complexes. The Journal of biological chemistry 283, 1692–1704, doi:10.1074/jbc.M707257200 (2008).

80 Hervera, A. et al. Reactive oxygen species regulate axonal regeneration through the release of exosomal NADPH oxidase 2 complexes into injured axons. Nat Cell Biol 20, 307–319, doi:10.1038/s41556-018-0039-x (2018).

81 Benedikter, B. J. et al. Redox-dependent thiol modifications: implications for the release of extracellular vesicles. Cell Mol Life Sci 75, 2321–2337, doi:10.1007/s00018-018-2806-z (2018).

82 Straube, A. & Merdes, A. EB3 regulates microtubule dynamics at the cell cortex and is required for myoblast elongation and fusion. Curr Biol 17, 1318–1325, doi:10.1016/j.cub.2007.06.058 (2007).

83 Eisenhoffer, G. T. et al. Crowding induces live cell extrusion to maintain homeostatic cell numbers in epithelia. Nature 484, 546–549, doi:10.1038/nature10999 (2012).

