## Supplemental data files (Figures S1-S10, Movie legends 1-14, Materials & Methods) for "“Epidermal Eg5 promotes X-ROS dependent paclitaxel neurotoxicity”"

**Affiliations:**
[1]Department of Biology, University of Miami, Coral Gables, Florida 33146, USA
[2]Department of Cell Biology, University of Miami Miller School of Medicine, Miami, Florida 33136, USA
[3]University of New England, Department of Biomedical Sciences, Biddeford, Maine, 04005
[4]Department of Computer Science, University of Miami, Coral Gables, Florida 33146, USA
[5]MDI Biological Laboratory, Kathryn W. Davis Center for Regenerative Biology and Medicine, Bar Harbor,
[6]Department of Chemical and Biomedical Engineering, University of Maine, Orono, Maine 04469, USA [7]Department of Neurology, Mayo Clinic, Rochester, MN 55905, USA
Maine 04762, USA
[8]Sylvester Comprehensive Cancer Center, University of Miami Miller School of Medicine, Miami, Florida 33136, USA

### Supplemental Figures & Legends

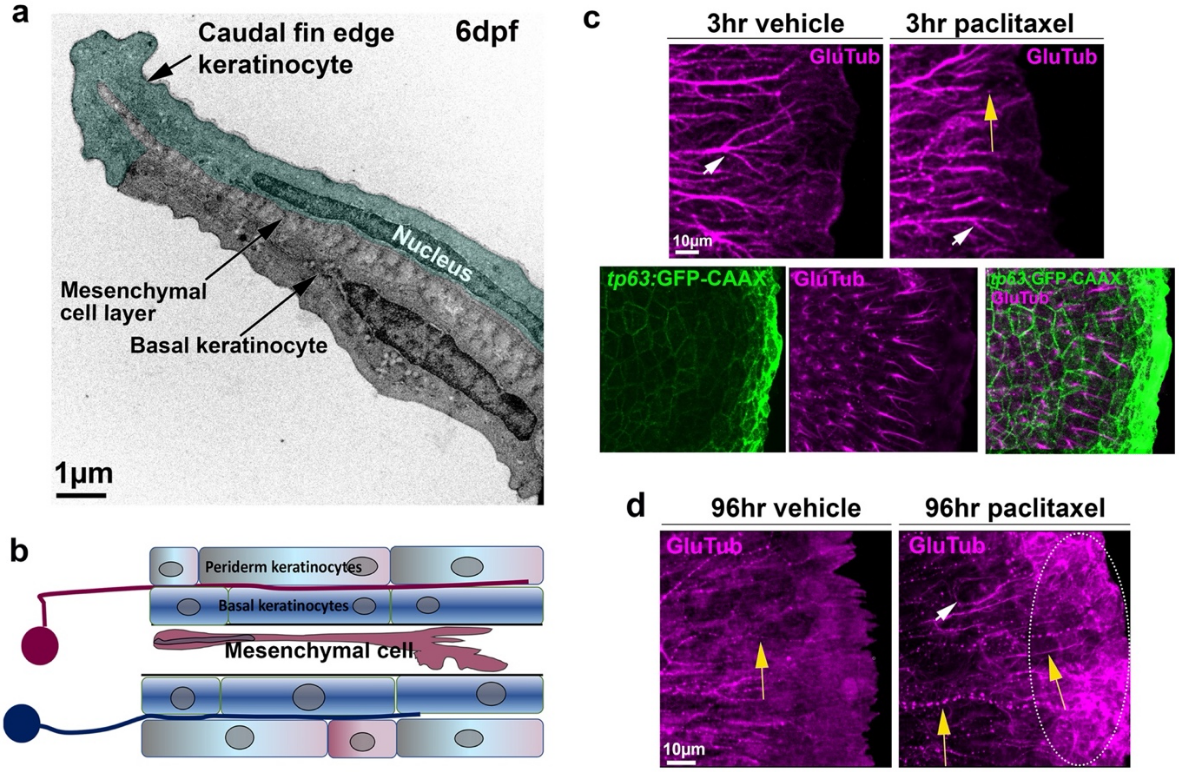


**Figure S1. Detyrosination in the zebrafish caudal fin.** (**a**) Transmission electron microscopy of a larval zebrafish caudal fin at 6 days post fertilization (dpf) showing an infolded epidermis and a single keratinocyte wrapping around the distal caudal fin edge (green). Mesenchymal cells are located medially between the infolded epidermis. (**b**) Schematic of the caudal fin shown in (**a**) to depict the cell types that are present. Two contralateral sensory neurons (red and blue) are shown that innervate the epidermis and arborize between the periderm and basal keratinocyte layer. (**c**) GluTub staining of detyrosinated (stabilized) microtubules (dMTs) in a 2dpf zebrafish embryo reveals strong labelling in mesenchymal cells (arrows) of both 3hr vehicle (0.05% DMSO) and paclitaxel treated fish. Axonal dMTs are evident following paclitaxel treatment (yellow arrow) but not present in vehicle controls. Keratinocyte plasma membranes are visualized with *tp63*:GFP-CAAX (green) to demonstrate that mesenchymal cell microtubules span multiple keratinocyte cell diameters. (**d**) At 6dpf, dfMTs are present in mesenchymal cells and axons of vehicle-treated fish. Following 96hr paclitaxel treatment, dMTs that are fasciculated form in individual keratinocytes in the caudal fin (white arrow) and most prominently at the fin edge (oval dotted line). Cutaneous axons (yellow arrows) display a weak punctate detyrosination pattern in the control animal, which is more prominent following paclitaxel treatment.

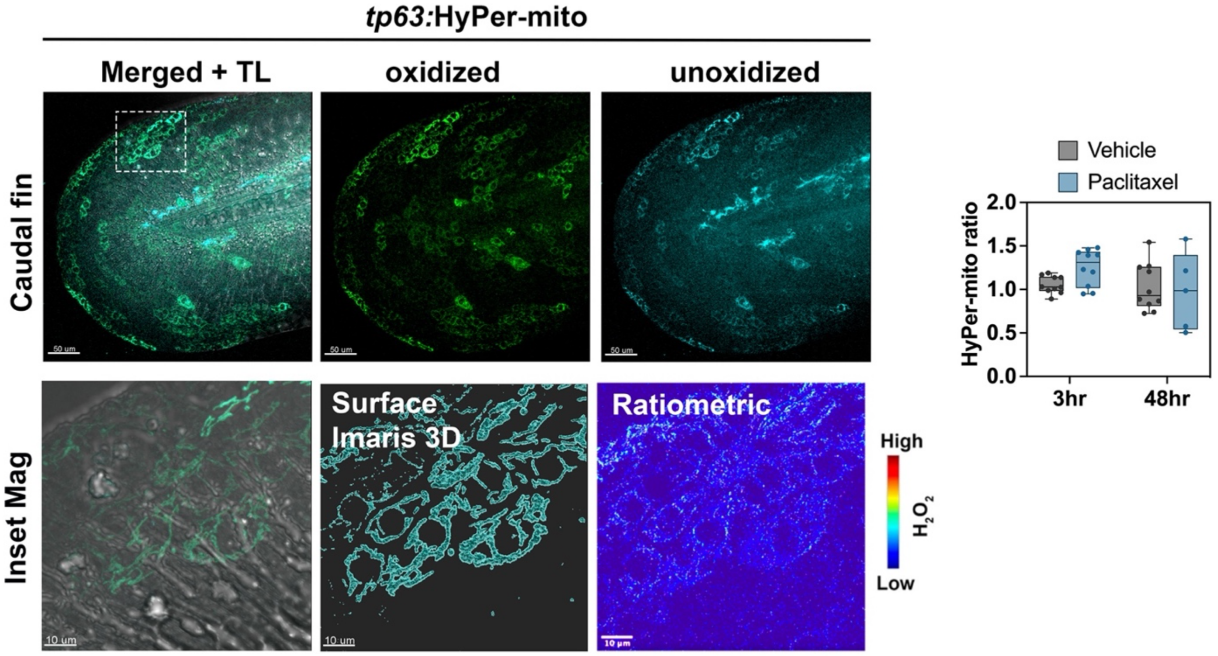


**Figure S2. Keratinocyte mitochondria show subtle changes following paclitaxel treatment.** Mitochondria in basal keratinocytes are labelled with *tp63*:HyPer-mito. Ratiometric imaging shows that oxidation is not significantly different when comparing 3 and 48hr vehicle versus 22μM paclitaxel treatment. HyPer-mito ratio: 505/420nm.

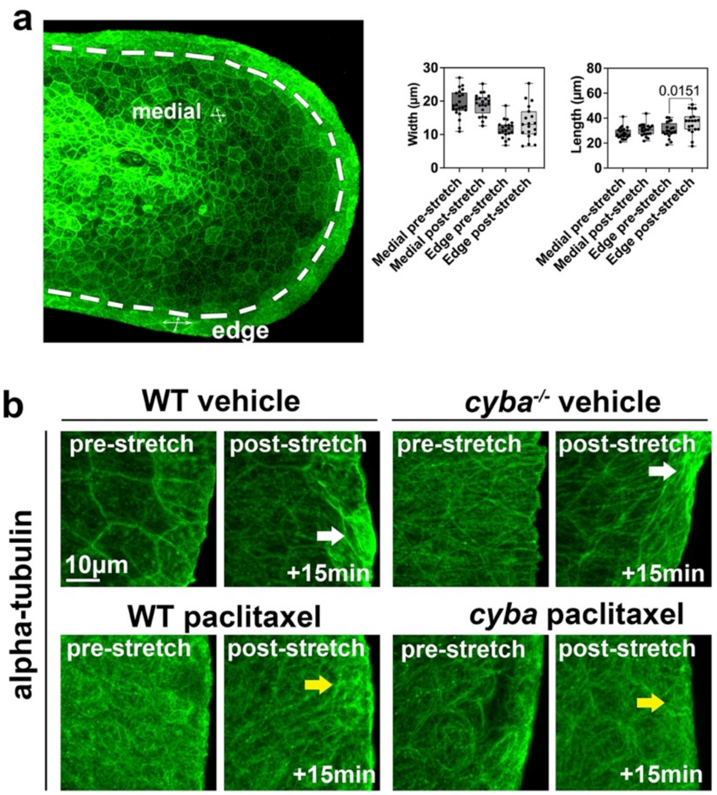


**Figure S3. ZStretcher validation in larval zebrafish.** (**a**) Tg(*tp63*:GFP-CAAX) transgenic animals with fluorescently labelled basal keratinocyte plasma membranes were used to measure keratinocyte length and width by comparing medial and edge (dashed line) keratinocytes. Quantifications were performed before and after zebrafish stretch on the ZStretcher, which reveals that fin edge keratinocytes increase in length but not in width whereas no effect is seen for medial keratinocytes (n=4 animals). (**b**) Immunofluorescence staining for alpha tubulin shows microtubule stretching along the fin edge in wildtype and homozygous *cyba*$^{-/-}$ fish treated for 48 hours with vehicle whereas microtubule stretching at the fin edge is less evident following paclitaxel treatment (white versus yellow arrows).

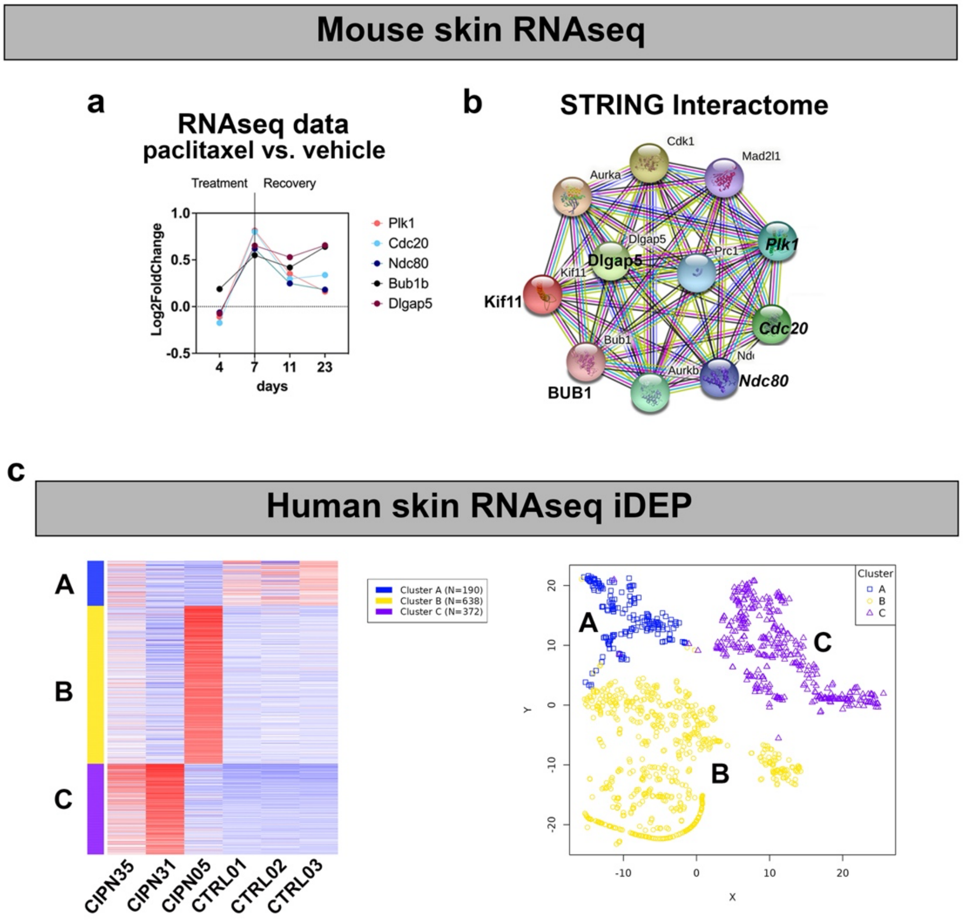


**Cluster A**

| Cluster | adj.Pval | nGenes | Pathways |
|---|---|---|---|
| A | 3.7e-06 | 10 | Keratinization |
| | 4.9e-06 | 15 | Epithelial cell differentiation |
| | 1.5e-05 | 10 | Keratinocyte differentiation |
| | 1.5e-05 | 7 | Cornification |
| | 3.7e-05 | 6 | Gluconeogenesis |
| | 3.7e-05 | 10 | Epidermal cell differentiation |
| | 3.7e-05 | 6 | Hexose biosynthetic process |
| | 4.7e-05 | 6 | Monosaccharide biosynthetic process |
| | 7.7e-05 | 10 | Skin development |
| | 1.0e-04 | 9 | Lipid catabolic process |
| | 1.1e-04 | 20 | Tissue development |
| | 1.1e-04 | 16 | Epithelium development |
| | 1.5e-04 | 10 | Epidermis development |
| | 2.9e-04 | 16 | Lipid metabolic process |
| | 4.6e-04 | 3 | One-carbon compound transport |

**Cluster B 5 weeks**

| | | | |
|---|---|---|---|
| B | 4.2e-07 | 5 | Cornification |
| | 4.3e-06 | 3 | Oxygen transport |
| | 5.6e-06 | 3 | Gas transport |
| | 5.6e-06 | 5 | Keratinization |
| | 7.3e-06 | 9 | Cell death |
| | 9.9e-06 | 5 | Keratinocyte differentiation |
| | 9.9e-06 | 3 | Hydrogen peroxide catabolic process |
| | 1.8e-05 | 3 | Bicarbonate transport |
| | 1.9e-05 | 5 | Epidermal cell differentiation |
| | 3.0e-05 | 5 | Skin development |
| | 4.4e-05 | 5 | Epidermis development |
| | 2.0e-04 | 3 | Cellular oxidant detoxification |
| | 3.0e-04 | 3 | Cellular response to toxic substance |
| | 3.2e-04 | 5 | Epithelial cell differentiation |
| | 3.2e-04 | 3 | Response to hydrogen peroxide |

**Cluster C 31/35 weeks**

| | | | |
|---|---|---|---|
| C | 8.3e-06 | 7 | Chromatin assembly or disassembly |
| | 1.1e-05 | 7 | DNA packaging |
| | 2.2e-05 | 7 | Chromatin remodeling |
| | 2.2e-05 | 6 | Chromatin assembly |
| | 3.9e-05 | 6 | Nucleosome organization |
| | 6.0e-05 | 7 | DNA conformation change |
| | 8.4e-05 | 7 | Protein-DNA complex subunit organization |
| | 1.3e-04 | 5 | Nucleosome assembly |
| | 4.0e-04 | 6 | Protein-DNA complex assembly |
| | 9.0e-04 | 7 | Chromatin organization |
| | 3.9e-03 | 3 | Chromosome condensation |
| | 5.2e-03 | 8 | Chromosome organization |

**Figure S4. Paclitaxel treatment induces cell cycle regulators in the skin.** (**a**) Validation of mouse RNAseq data confirms the upregulation of spindle checkpoint regulators (*Plk1m Cdc20, Ndc80, Bub1, Dlgap5*) following paclitaxel treatment relative to vehicle controls, with the highest expression increase at D7 during peak neuropathy. (**b**) STRING interactome analysis predicts the interactions of these genes with *Kif11* and other mitotic genes. (**c**) iDEP.96 analysis of the 1200 most variable differentially expressed genes shows three distinct clusters. The first cluster (A, blue) harbours genes that are downregulated in the CIPN patients. Cluster B (yellow) harbours genes that are upregulated in the patient diagnosed with CIPN 5 weeks prior. Cluster C (purple) harbours genes that are upregulated in the patients diagnosed with CIPN 35 and 31 weeks prior).

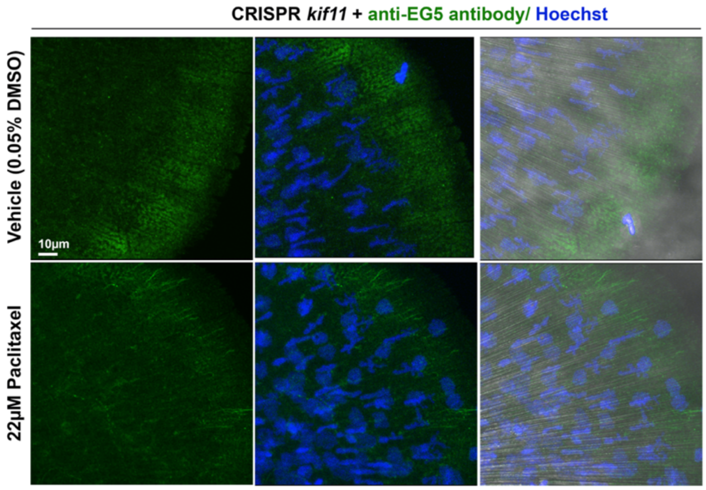


**Figure S5. Zebrafish *kif11* CRISPR knockout without EG5 induction in caudal fin following paclitaxel treatment.** Vehicle (top) and paclitaxel (bottom) treatment for 96 hours does not induce Eg5 expression detected with an Eg5 specific antibody using immunofluorescence staining.

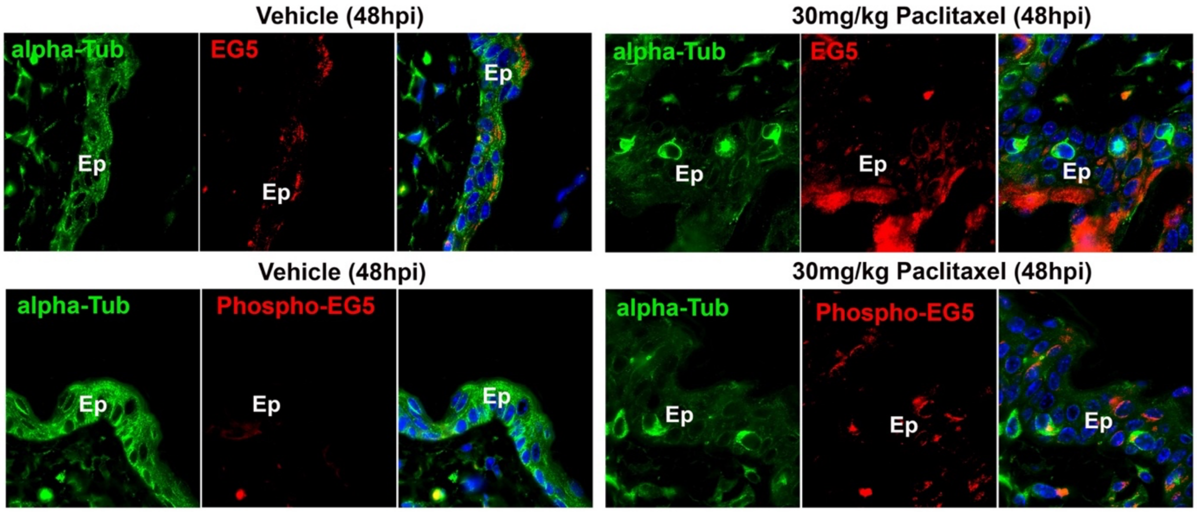


**Figure S6. Paclitaxel induces total and phospho-Eg5 in mouse suprabasal keratinocytes.** Mice were i.p. injected once with either 0.9% NaCl (vehicle) or 30mg/kg paclitaxel, followed by fixation 48 hours post injection. Alpha-tubulin antibody staining is used as counterstain. DAPI is used to depict nuclei. Top panel: Weak EG5 expression in the vehicle control epidermis (left) but strong induction of EG5 following paclitaxel treatment (right). Lower panel: Eg5 phosphorylation is absent in the skin following vehicle injection (left). Paclitaxel injection induces Eg5 phosphorylation in basal and suprabasal keratinocytes. *Ep: Epidermis; hpi: hours post injection*

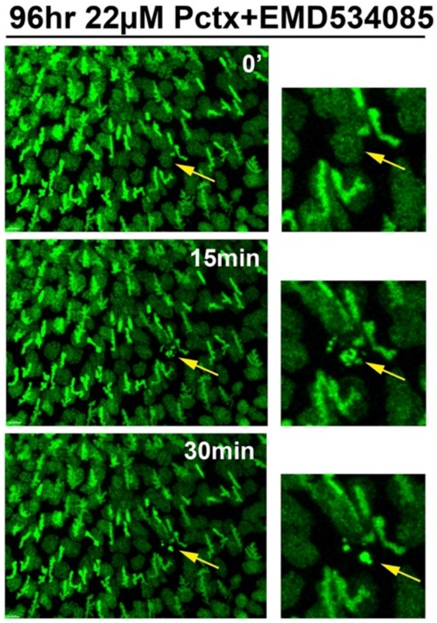


**Figure S7. Time-lapse imaging of keratinocyte cell death in caudal fin.** Caudal fin keratinocyte nuclei are visualized in a Tg(*h2a*:h2a-GFP) zebrafish caudal fin at 6dpf. The fish was treated with paclitaxel+EMD534085 for 96hr. The intact keratinocyte nucleus (top, yellow arrow) condenses within 15 minutes (middle, yellow arrow) and subsequently fragments (bottom, yellow arrow), indicative of cell death. Right panels show a higher magnification images.

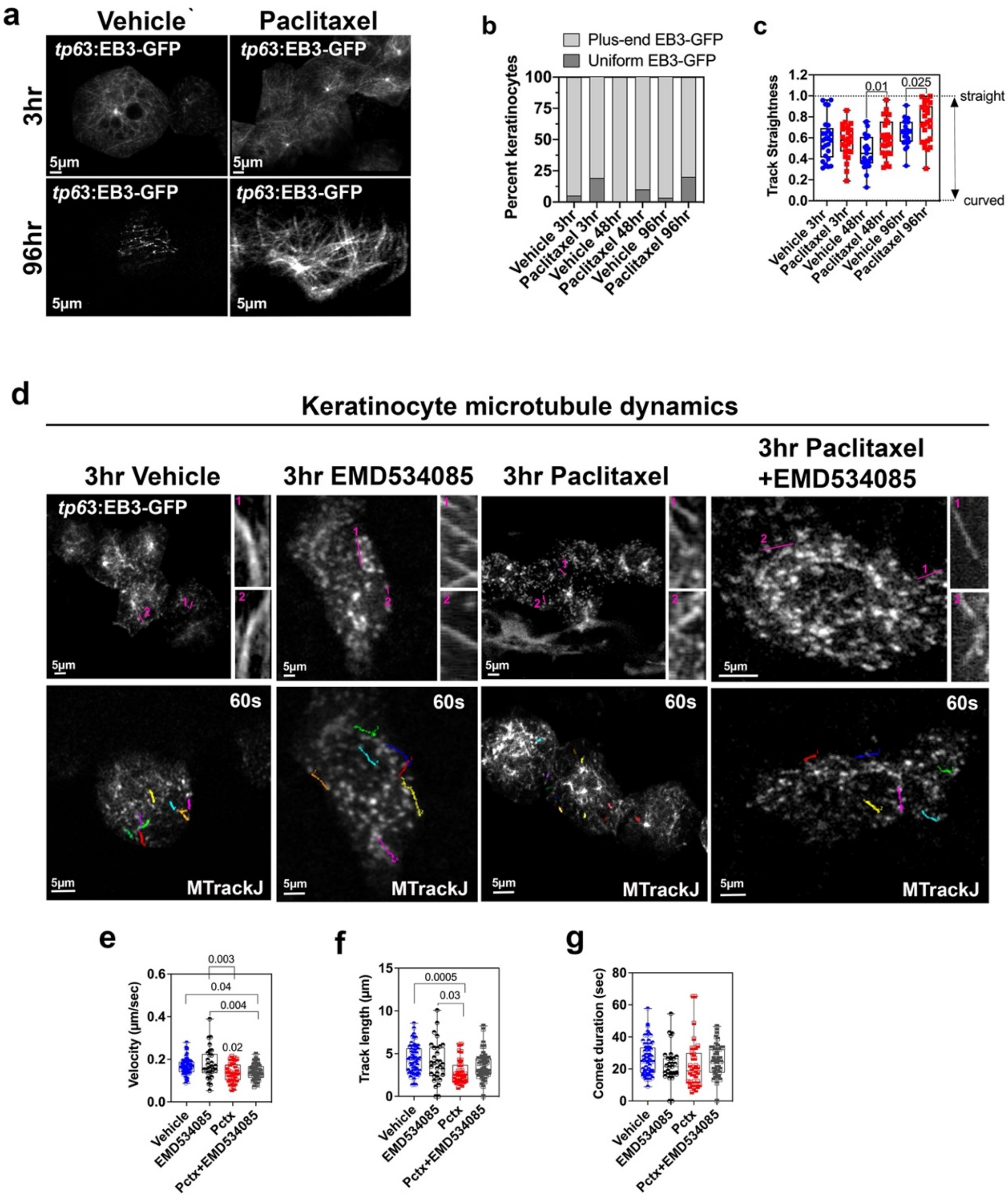

**Figure S8. Paclitaxel modulates microtubule growth dynamics in keratinocytes independent of Eg5.** Microtubule growth dynamics were captured and quantified in caudal fin keratinocytes expressing *tp63*:EB3-GFP, see also Movies 10 and 11. (**a, b**) Keratinocytes with uniform EB3-GFP localization along the microtubule lattice. Paclitaxel treatment increases the number of keratinocytes with uniform EB3-GFP labelling (n=5 animals). (**c**) Increased straightness of microtubule growth tracks following 48 and 96hr paclitaxel treatment (1=straight, 0=curved). (**d**) Kymographs of growing microtubules generated with MTrackJ shows more rapid depolymerization in the presence of paclitaxel, consistent with quantifications in f. (**e**) Significantly decreased comet velocity following 3hr paclitaxel treatment is not rescued by co-administration of EMD534085 (n≥8 animals/group). (**f**) Significantly decreased track length upon 3hr paclitaxel treatment is not rescued by co-administration of EMD534085 (n≥8 animals/group). (**g**) Comet duration is not significantly changed for any of the treatments (n≥8 animals/group).

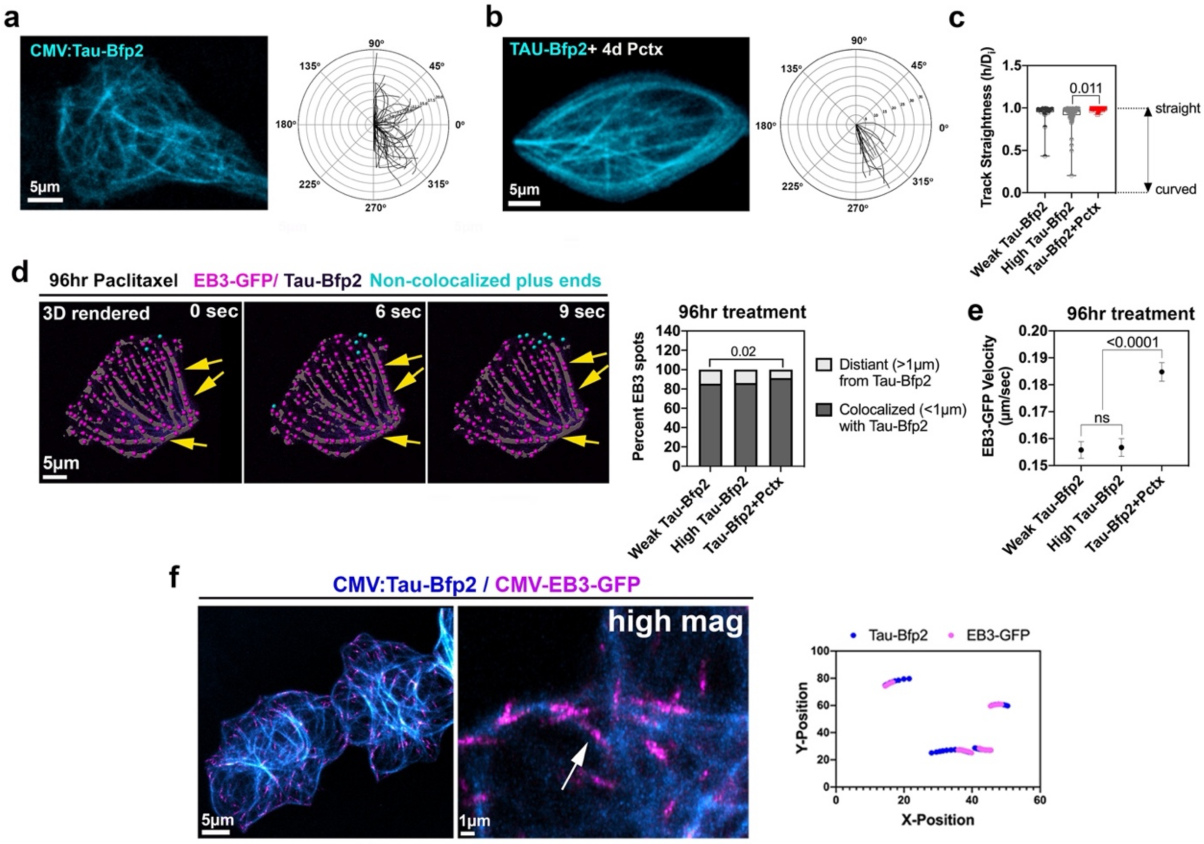


**Figure S9. Microtubule behaviour in genetically stabilized microtubules.** (**a**) Microtubule network in *CMV*-Tau-Bfp2 expressing keratinocytes following 96hr vehicle treatment. Polar plot shows tracings to outline microtubule conformations (n=5 animals/ plot). (**b**) 96hr paclitaxel treatment induces linearization of Tau-Bfp2-stabilized microtubules. The polar plot validates the increased linearity (n≥4 animals/ plot). (**c**) Microtubule track straightness in keratinocytes expressing Tau-Bfp2 with and without 96hr paclitaxel treatment shows enhanced linearization in the presence of paclitaxel: Straightness (1= straight, 0=curved) (n≥5 animals/group). (**d**) 3D reconstruction of EB3-GFP plus-ends in a keratinocyte following 96hr paclitaxel treatment. More than 80% of EB3-GFP plus-ends colocalize with Tau-Bfp2 regardless of low or high Tau expression. Paclitaxel treatment for 96hr further increases the colocalization (n≥5 animals/group). (**e**) Paclitaxel treatment for 96hr increases EB3-GFP velocity in the presence of Tau-Bfp2. (**f**) Colocalized EB3-GFP and Tau-Bfp2 microtubules (arrow) show overlapping growth tracks when traced along curvatures.

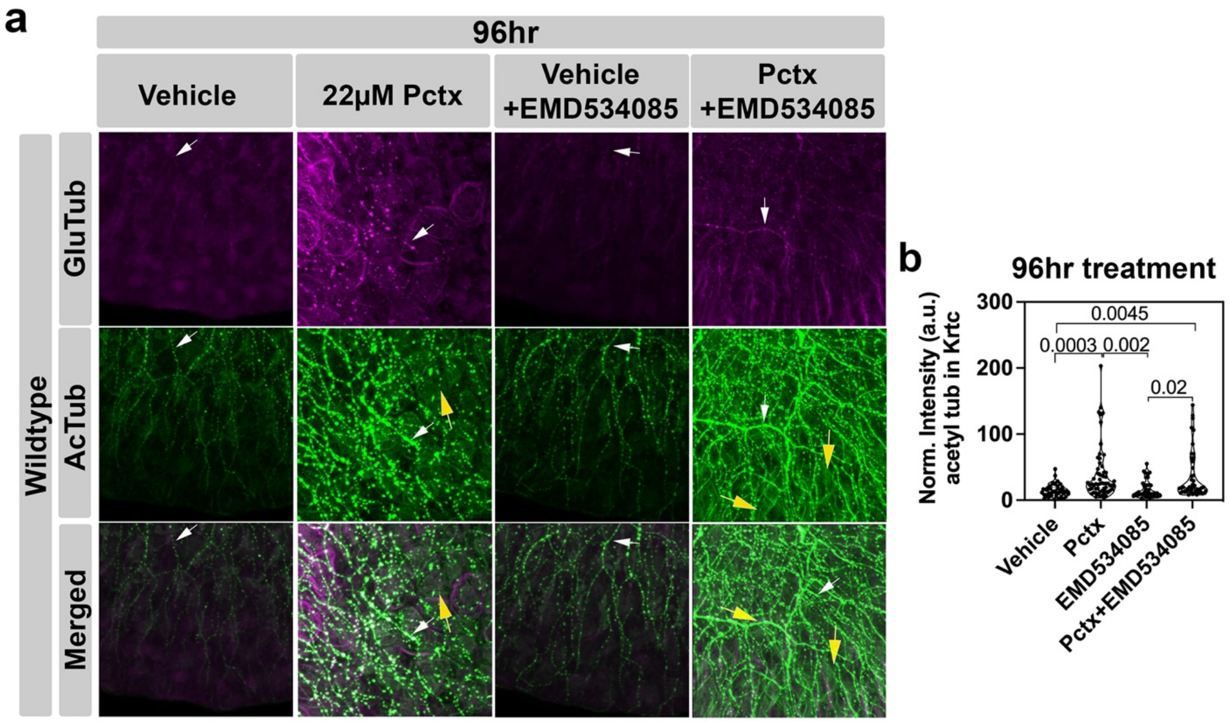


**Figure S10. Paclitaxel induces axonal detyrosination and keratinocytes acetylation.** Increased axonal dMT formation (white arrows) and keratinocyte-specific microtubule acetylation at K40 (yellow arrows) following 22μM paclitaxel treatment for 96 hours with and without EMD534085. (**b**) Quantification of normalized acetylation intensity shows a significant increase in keratinocyte microtubule acetylation after 96hr paclitaxel treatment with and without EMD534085.

**Supplemental Movies Legends**

**Movie S1. Mesenchymal cell morphology.** 3D reconstruction of detyrosinated microtubules (magenta) in a mesenchymal cell at 3 days post fertilization. The nucleus is shown in blue.

**Movie S2. Detyrosinated microtubules (anti-GluTub) and nuclei (Hoechst 33342) in the caudal fin of a 96hr vehicle-treated zebrafish (6dpf)**. Detyrosination is primarily present in mesenchymal cells that appear as elongate structures crossing multiple keratinocytes. Mesenchymal cell nuclei are also elongated and jagged, unlike the rounded keratinocyte nuclei.

**Movie S3. Detyrosinated microtubules detected with anti-GluTub antibody staining and nuclei detected with Hoechst 33342 staining in the caudal fin of a 96hr paclitaxel-treated zebrafish (6dpf).** Keratinocyte-specific dfMTs wrapping around nuclei.

**Movie S4. 3D reconstruction of Nox1 staining in a single epidermal keratinocyte following vehicle treatment**. Shown is a 7dpf zebrafish following 120hr treatment with 0.05% DMSO. Nox1 staining is punctate within the cytoplasm and nucleus and uniformly locates at the plasma membrane.

**Movie S5. 3D reconstruction of Nox1 staining in a single epidermal keratinocyte following paclitaxel treatment.** 3D rendered Nox1 staining in a single epidermal keratinocyte of a 7dpf zebrafish following 120hr paclitaxel treatment (22μM). Nox1 appears clustered inside the nucleus and in cytoplasmic vesicles, and uniformly locates at the plasma membrane.

**Movie S6. 3D reconstruction of a nucleus and surrounding dfMTs in a caudal fin keratinocyte following paclitaxel treatment.** The dfMTs wrap around and partially associate with the nucleus following 96hr paclitaxel treatment.

**Movie S7. 3D reconstruction of dfMTs pinching off nuclear content.** 3D reconstructed dfMTs appear to pinch off nuclear content following 96hr paclitaxel treatment.

**Movie S8. Fluorescence staining and 3D reconstruction of a nucleus that is perforated by dfMTs.** Shown are two movie sequences of a nucleus that is perforated by dfMTs following 96hr paclitaxel treatment. The first sequence shows GluTub fluorescence staining to label dfMTs and Hoechst33342 staining labelling the nucleus. The second sequence shows a 3D reconstruction of the dfMTs and nucleus.

**Movie S9. Live imaging of cell division.** Cell divisions (blue arrows) are visible around the notochord and spinal cord region in a 6dpf transgenic Tg(*h2a*:h2a-GFP) zebrafish larva.

**Movie S10. Plus-end tracking of keratinocyte microtubules in a vehicle-treated zebrafish.** Transiently injected *tp63*:EB3-GFP labels growing plus-ends of keratinocyte microtubules following 96hr vehicle treatment. EB3-GFP was traced for 60sec at 1 frame/sec using MTrackJ (Fiji).

**Movie S11. EB3 is uniformly distributed along microtubules following paclitaxel treatment.** Transiently injected *tp63*:EB3-GFP is uniformly distributed along keratinocyte microtubules following 96hr paclitaxel (22μM) treatment. EB3-GFP was traced at 1 frame/sec using MTrackJ (Fiji).

**Movie S12. Co-expression of *tp63*:EB3-GFP and *CMV*:Tau-Bfp2 in a vehicle-treated fish.** *tp63*:EB3-GFP (magenta) and *CMV*:Tau-Bfp2 (blue) co-expression in a basal keratinocyte following 96hr vehicle (0.05% DMSO) treatment shows dynamic microtubule growth.

**Movie S13. Co-expression of *tp63*:EB3-GFP and *CMV*:Tau-Bfp2 in a paclitaxel-treated fish.** *tp63*:EB3-GFP and *CMV*:Tau-Bfp2 co-expression in a basal keratinocyte following 96hr paclitaxel (22µM) treatment shows reduced and linear microtubule growth along pre-existing Tau-stabilized microtubules.

**Movie S14. 3D reconstruction of EB3-GFP and Tau-Bfp2 following paclitaxel treatment.** 3D reconstruction of microtubule plus-ends growing along pre-existing Tau-stabilized microtubules as shown in **Movie S13.** Microtubule plus-ends were detected using the Spots function and Tau-Bfp2 labelled microtubules were detected using the Surface function in Imaris (Bitplane).

### Supplemental Materials and Methods

#### Zebrafish husbandry and transgenic lines

<u>Wildtype strains:</u> Zebrafish (Nacre/mitfa), Tuebingen and AB strains were purchased from the Zebrafish International Resource Center (ZIRC) and embryos raised and bred according to NIH guidelines. Tuebingen wildtype fish were used for CRISPR studies. Nacre fish were used for membrane labelling of keratinocytes in *tp63*:GFP-CAAX. All remaining analyses were performed in AB fish. Animals were handled in strict accordance with good animal practices as approved by the appropriate IACUC committees (MDI Biological Laboratory IACUC number #A13-20; University of Miami's #A-3224-01 and AALAC accreditation site: 001069). Zebrafish eggs were collected in a strainer and rinsed with deionized water, and then transferred into Petri dishes with Embryo medium (Instant Ocean salt water + methylene blue, Zebrafish book protocol) or Ringers solution. Following overnight incubation at 28.5°C, the embryos were cleaned, and fresh Ringer's solution with phenol-thio-urea added for further incubation. The embryos were kept in a 14:10hr light/dark cycle.
<u>Transgenic lines:</u> Tg(*h2a*:*h2a*-GFP, Cat. No. ZL1087) transgenic fish were obtained from the Zebrafish International Resource Center (ZIRC). *tp63*:GFP-CAAX fish were previously published[1].

#### Plasmids

<u>*CREST3:EB3-GFP:*</u> pCS2_CMV:EB3-GFP was a gift from the Koester Lab (University of Braunschweig, Germany). The CMV promoter was removed from pCS2 by digestion with HindIII, followed by Klenow fragment incubation to create blunt-ends, purification and subsequent digestion was done with SalI. The *CREST3* promoter in Tol2_*CREST3*:GFP (gift from Alvaro Sagasti, UCLA) was digested with EcoRV (created blunt-end site) and SalI. The gel-purified fragment was ligated into the pCS2_EB3-GFP plasmid overnight at 16°C and transformed into Top10 cells for purification and injection.
<u>*tp63:EB3-GFP*</u> was generated by removing the CMV promoter as above. The *tp63* promoter was removed by digestion of pBSK-JC_T2_*tp63*_Gal4VP16_GFP-5xUAS-MCS with ApaI and AvrII, followed by Klenow treatment to generate blunt ends. Following gel purification, the *tp63* promoter fragment was ligated into pCS2-EB3-GFP, as above. Both constructs were sequenced and further verified by *in vivo* imaging.

tp63:*kif11*-AcGFP was generated by PCR amplification of zebrafish *kif11* from cDNA (Horizon Discovery, Clone ID: 3815942) for ligation into the pAcGFP-N1 (Takara, Cat. No. 632501) fusion vector. The following primers were used: Fwd 5'-AAGGCCTCTGTCGACCATGGCATCATCACAAGTAC-3' and rev 5'-AGAATTCGCAAGCTTATTCTGACATCTGAGTGGAAGT-3' and Q5 Polymerase (NEBNext® High-Fidelity 2X PCR Master Mix). This was followed by PCR purification (QIAGEN PCR Purification kit) and ligation of the amplicon into pAcGFP-N1. The plasmid was transformed into One-Shot Top10 cells (ThermoScientific, Cat. No. C404010) and plasmids were purified from colony minipreps using the MiniPrep kit (QIAGEN). The insert (*kif11*-AcGFP) was subsequently amplified from using the following primers: Fwd 5'-GGATCCTTATGGCATCATCACAAGT-3' and Rev 5'-GCGGCCGCTTCTTGTACAGCTC-3' that added BamHI/NotI restriction sites. KXIG:*tp63*:AcGFP was digested with BamHI/NotI to remove AcGFP with gel extraction (QIAGEN) and BamHI_*kif11*-AcGFP_NotI was inserted via ligation using T4 DNA ligase (Promega, Cat. No M180A) at room temperature for 2 hours. The ligation reaction was transformed into One-Shot Top10 cells and colonies were screened via PCR for positive inserts. The positive colonies were prepped using the QIAGEN MiniPrep kit and verified via sequencing. Positive plasmids were injected into zebrafish and verified for fluorescence, which is only visible if *kif11* is cloned in-frame with AcGFP.

*isl1:Gal4VP16_14xUAS-tdTomato* was a gift from Alvaro Sagasti (UCLA).

*mTagBFP2-MAPTau-C-10* was provided by Michael Davidson (Addgene plasmid #55311; http://n2t.net/addgene:55311;RRID:Addgene_55311).

*tp63:HyPer-CAAX:* KXIG:*tp63*:AcGFP was digested with BamHI/NotI (NEB) to excise AcGFP. Hyper was amplified from pHyPer-dMito (Evrogen) without the stop codon using the following primers: Fwd: 5'-3' catttacctctgaagccacgggtttagtgaaccgtcag and Rev: 5'-ttcctcctccAACCGCCTGTTTTAAAAC-3'. The CAAX motif was amplified using the primers 5'-acaggcggttGGAGGAGGAAGATCTAAG-3' and 5'-tcgagctccaccgcggtggcAACACCCCTTGTATTACTG-3' from the pME-EGFP-CAAX vector (gift from Chi-Bin Chien). Both amplicons were purified using the PCR Purification kit (QIAGEN) and ligated via HiFi Assembly using the 2x HiFi Master mix (NEB, Cat. No. M0541) at 50ºC for 20 minutes. The assembly reaction was transformed into NEB 5-alpha competent cells provided with the HiFi Assembly kit (NEB, Cat. No. E5520) and colonies grown and purified via the Plasmid Miniprep kit (Qiagen) and subsequently sequenced.

*tp63:HyPer-mito:* KXIG:*tp63:HyPer-mito* was cloned by digesting *tp63*:AcGFP with BamHI/NotI (NEB) to excise AcGFP. Hyper-mito was amplified from pHyPer-dMito (Evrogen) using the following primers Fwd: 5'-catttacctctgaagccacgggtttagtgaaccgtcag-3' and Rev: 5'-tcgagctccaccgcggtggctaagatacattgatgagtttgg-3'. The amplicon was ligated into the vector containing the *tp63* promoter using the Hi-Fi Assembly Master mix (NEB) at 50ºC for 20 minutes. The assembly reaction was transformed into NEB 5-alpha competent cells and purified using the plasmid miniprep kit by Qiagen, and subsequently sequenced.

**Transmission Electron Microscopy.** TEM methods were published in [2].

#### Pharmacological agents

Paclitaxel was purchased from Sigma-Aldrich (Cat No. T7402) and upon arrival stored as powder at 4°C. A stock solution was subsequently prepared in 100% fresh DMSO to make

5.9mM paclitaxel using the entire bottle to avoid imprecise dilutions. The stock solution was then aliquoted into 20-30µl aliquots and stored at -20°C until use (maximal 6 months). We noticed that the solution is not as effective after 6 months. Immediately prior to use, paclitaxel was diluted in Ringer's solution and added to larval zebrafish that were dechorionated. The larval fish were placed individually or in small groups into wells of a 12-well plate containing the treatment solutions. 0.05% DMSO was used as control since we noticed that equal amounts of DMSO in comparison to treatment concentrations promote toxicity that is not observed when DMSO is used as solvent in combination with chemical compounds. The plates were returned to 28.5°C and protected from light. Treatment lengths are indicated in the text. The paclitaxel and DMSO solutions were exchanged every 48hr if longer incubations were used. EMD534085 - Eg5 inhibitor (MedChemExpress, Cat. No. HY-15000) was diluted in 100% DMSO upon arrival to make a 10mM stock solution and stored at -20°C. Immediately prior to use, the inhibitor was diluted to 25µM and added at 4dpf, following two days of either vehicle or paclitaxel incubation. For Nox1 staining experiments, the inhibitor was added together with paclitaxel at 2dpf. The solutions were exchanged every 48hr.

**Immunofluorescence**

Zebrafish immunofluorescence staining: Larval fish were transferred into 20ml glass vials, followed by fixation in 4% paraformaldehyde (PFA)/1x Phosphate-buffered saline (PBS) for 1.5hr at room temperature, gently rocking. Following a 5-minute incubation in 1xPBS+0.1% Tween-20 (PBST), rocking at room temperature, zebrafish were permeabilized in 1xPBS+0.1% Triton X-100 for 10-30min, also rocking at room temperature. This was followed by transfer of larval fish into 2ml reaction tubes and incubation in blocking buffer (1xPBST+5%BSA) for 30min at room temperature, rocking. Larval fish were incubated overnight in antibody solution (Millipore Sigma, Cat. No. AB3201, 1:300 rabbit anti-GluTub; Sigma, Cat No. T6793, 1:500 mouse anti-acetylated tubulin) in blocking buffer at 4°C, on a rotator. The next morning, larval fish were transferred into 6-well plates and washed 4x15min in 1xPBST at room temperature, rocking. This was followed by incubation in secondary antibody (goat anti-rabbit Cy5, Abcam, Cat. No. ab97077; goat anti-mouse Cy3, Abcam, Cat. No. ab97035) and 1:10,000 Hoechst 33342 (ThermoFisher Scientific, Cat. No. 62249, 20mM solution) in blocking buffer for 1hr at room temperature, rocking. Fish were covered to avoid bleaching. The fish were washed 4x15min in 1xPBST at room temperature, rocking, and immediately mounted for imaging on glass bottom petri dishes (Spectrum Laboratory Products, Cat. No. 750-10403-UE) using 1% agarose (Thermo Fisher Scientific, Cat. No. 16520050). A 1:200 dilution was used for mouse alpha-tubulin antibody (Proteintech, Cat. No. 66031-1) and a 1:1,000 dilution of donkey anti-mouse IgG Alexa488 (ThermoFisher Scientific, Cat. No. A-21202) for secondary detection. For Nox1 immunostaining, a 1:300 dilution of primary antibody (Anti-NOX1 Rabbit Polyclonal Antibody, Avantar, Cat. No. 102164-796) and a 1:1,000 dilution of secondary goat anti-rabbit Cy2 (Abcam, Cat. No. ab6940) antibody was used. Eg5 antibody (Abcam, Cat. No. 61199) was used at a 1:100 and secondary goat anti-rabbit Cy2 (Abcam, Cat. No. ab6940) at 1:1,000 dilution.

**Quantitative PCR**

Zebrafish larvae (AB/Nacre) were treated in pools of ~20-30 larvae per treatment per biological replicate (BR); each BR consisted of embryos from distinct parents. Larvae were collected into 1.5ml reaction tubes post-treatment and RNA extraction was performed using the RNeasy Plus Micro kit (Qiagen, Cat. No. 74034). cDNA was synthesized from ~300ng total RNA using the Superscript IV VILO kit (Thermo Fisher, Cat. No. 11756050). Primers in target genes were designed such that the primers annealed in two exons separated by one or several large introns that could not be amplified with the selected PCR settings, to avoid genomic DNA contamination. Quantitative PCR was performed with the Applied Biosystems QuantStudio 3 using PowerUp SYBR Green (ThermoFisher, Cat No. A25741). Amplification signals for expressed target genes were normalized to zebrafish 18s rRNA signals. qPCR conditions were used as follows: 95ºC for 10 minutes, 40 cycles of 95ºC for 15 seconds, 50ºC for 30 seconds, 72ºC for 30 seconds. Each biological replicate was run in quadruplicates. Data is presented as relative expression compared to control using the delta-delta Ct (2^-ΔΔCt) method.

Primer sequences for qPCR:

| **Gene** | **Primer** | **Sequence (5' → 3')** | **Target Exon** |
|---|---|---|---|
| *18s* | forward | ATGTCCCTCGTCATCCCAGAGAAGTT | 2 |
| *18s* | reverse | ATTGTCCAGACCATTAGCAAGGA | 5 |
| *duox* | forward | TTCTTGGTCTGCCTTTGACG | 20 |
| *duox* | reverse | GCATGGAAGTACTTGGTAAG | 21 |
| *mmp13a* | forward | AATTACCTGACTCGACTGTATGG | 2 |
| *mmp13a* | reverse | CCAGTGGCGAAGAAGATCA | 3 |
| *nox1* | forward | TGGCAATAAACATCGCTTTG | 2 |
| *nox1* | reverse | TTCACTGGAGTCTTGGAGCA | 5 |
| *nox2* | forward | CGTATGTGCTCTCTCAGATTGG | 5 |
| *nox2* | reverse | GTCAATCCTGCTACTGTCGTAAA | 6 |

### ***kif11*** **CRISPR knockout**

CRISPR oligos targeting *kif11* were designed using the IDT CRISPR design tool (https://www.idtdna.com/site/order/designtool/index/CRISPR_CUSTOM). The oligo (5'-AGGTGACCGATCACCCAATG) was designed to anneal within exon 5 of 23 exons total in zebrafish *kif11* with expected mutations ~ position 270bp in the sequenced region using the following primers for PCR amplification: Fwd 5'- TTAGGTTTTTGGCCCTTCTG-3' and Rev 5'-GAGGGTCCTGATAGAGAAAAAGTGAA-3'. The forward primer was used for sequencing and yielded deletions in transiently injected embryos, as shown in example below in which single zebrafish were sequenced.

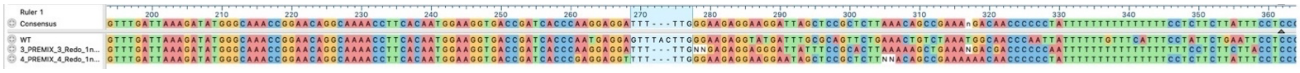

The CRISPR oligo was injected using the Alt-R system (IDT, https://www.idtdna.com/pages/technology/crispr/crispr-genome-editing/Alt-R-systems/cas9), which has provided highly reliable and efficient zebrafish knockout results in our lab. The IDT-recommended protocol (below) was used for CRISPR oligo preparations. Fertilized eggs were injected ~15 minutes post fertilization with CRISPR oligos to ensure maximal efficiency.

#### <u>Zebrafish embryo microinjection protocol (IDT)</u>

Ribonucleoprotein delivery using the Alt-R™ CRISPR-Cas9 System (Contributed by Jeffrey Essner, PhD, Associate Professor, Department of Genetics, Development, and Cell Biology, Iowa State University, Ames, IA, USA).

<u>Methods</u>

1. Resuspend Alt-R crRNA and tracrRNA in Nuclease-Free IDTE Buffer to final concentrations of 100μM each.
2. Mix the following components to create a 3μM gRNA solution:

| Component | Amount |
|---|---|
| 100µM Alt-R™ CRISPR-Cas9 crRNA | 3µL |
| 100µM Alt-R™ CRISPR-Cas9 tracrRNA | 3µL |
| Nuclease-Free Duplex Buffer (IDT) | 94µL |
| Final volume | 100µL |

3. Heat at 95ºC for 5 min.
4. Remove from heat and allow to cool to room temperature (15–25ºC) on bench top.
Note: the final concentration for the crRNA is 36ng/µL and for the tracrRNA is 67ng/µL.
5. Dilute Cas9 protein to a working concentration of 0.5µg/µL:

| Component | Amount |
|---|---|
| 10µg/µL Cas9 protein | 0.5µL |
| Cas9 working buffer (20 mM HEPES; 150 mM KCl, pH 7.5) | 9.5µL |
| Final volume | 10µL |

6. Assemble the RNP complexes, for each injection:
a. Combine 3µL of gRNA (from step 4) with 3µL of diluted Cas9 protein (from step 5).
b. Incubate at 37°C for 10 min.
c. Allow to cool to room temperature.

7. Collect embryos at the 1-cell stage and inject 3nL of RNP complex (from step 6).
Note: Dr Essner typically injects at least 15 embryos for each target and includes 5 uninjected embryos as controls.
8. Check injected fish for obvious toxicity at the following time points after fertilization: 8hr, 1, 2, 4 days
9. Isolate genomic DNA at day 4 after injection using the NaOH method.
Note: Pool 5 fish into 1 tube. Include 3 tubes for each injection and 1 tube of uninjected fish for a control.
10. Run PCR specific for your targeted region.
11. Analyse PCR products on 2% agarose gels to estimate mutation efficiency.
Note: Alternatively, sequence bands to determine accurate mutation frequency, or use a T7EI mismatch cleavage assay to estimate mutation efficiency.

**Imaging**

Immunofluorescence imaging of zebrafish was performed on the following confocal microscopes: Olympus FV1000, Zeiss LSM510, and LSM880 Airyscan. Each was used with a 20x air objective zooms between 1x-3.8x. For microtubule detection, 1µm sections in varying stack sizes were recorded to capture the full diameter of the caudal fin at either 512x512 (Zeiss LSM510), 860x860 (Olympus) or 1024x1024 (Zeiss LSM880) pixel resolution. Scanning was performed at variable scan speed to ensure high quality images. In general, slow scan speed was used for publication images and faster scan speed and the lower resolution was used for quantification purposes.
Live imaging of zebrafish: Fish were anesthetized in 2-phenoxyethanol (1:1000) in Ringer's solution prior to imaging and mounted in 1.2% agarose. For imaging of EB3-GFP and Tau-Bfp2, the same microscopes as above were utilized. The scan speed was kept at the maximum level in single slice mode with 1s intervals for a total of 60s to capture microtubule dynamics. *h2a*:H2A-GFP time-lapse recordings were produced using the Zeiss LSM880 AiryScan confocal microscope with 1µm sections recorded every 10min for 12hr using 800x800 pixels. All other live fish were imaged using 1024x1024 pixels on the Zeiss LSM880 AiryScan microscope.

Image processing: Images were processed and analysed in Imaris 9.5.1 (Bitplane, Switzerland) or Fiji. For publication images, projected stacks were saved as .tif files using the Snap tool, followed by processing in Photoshop to assemble the figures. 3D reconstructions were performed in Imaris using the semi-automated Surfaces tool, which includes background subtraction in addition to other features that allow for optimal background:noise detection. For example, to render nuclei, a region of interest was set in the caudal fin, followed by automated background subtraction, and object size adjustment. The surface details were set either automatically or manually adjusted depending on the final render quality. Threshold detection and voxel number settings were automatically set. The rendered images were adjusted through repeated re-adjustments of each setting, until a suitable fit was observed. The settings were kept the same for groups that were compared. Graphs were generated and statistics performed in Prism 9 (GraphPad) software. Schematics were prepared in Adobe Illustrator. HyPer mitochondria were ratiometrically displayed using the Image calculator function in ZEN Black whereby the 420nm unoxided and 505nm oxidized channels were added into the division calculator and the signal was amplified by a factor of 150.

**Quantifications**

Microtubule analyses were performed in Imaris 9.5.1 (Bitplane) by measuring the straightness of filaments (microtubules) using the "Filaments - Dendrite Straightness" tool, which is defined as the ratio between filament length and radial distance between two branch points (h). The value is always smaller than 1 since the Dendrite Straightness of 1 defines straight objects.
Fluorescence intensities were measured using the Imaris MATLAB plugin or Fiji. A line was typically placed over the region of interest and the fluorescence automatically quantified. The values were normalized to nuclear (Hoechst33342) fluorescence intensities within the same fish and region to obtain an intensity ratio.
Cell counting in the caudal fin was performed in three 100µm$^2$ boxes in the dorsal, medial and ventral caudal fin (~100µm from the edge).
Nuclear sphericity and volume were calculated in Imaris following 3D rendering of round keratinocyte nuclei using the Surface tool. First, background subtractions were performed, followed by automatic threshold detection. The threshold was adjusted if necessary to fit the nucleus. Thresholds were kept constant for comparisons.
Cell divisions and cell death were manually counted in 12hr movie recordings. The caudal fin was divided into 6 quadrants and individual cells in each quadrant were manually followed over 12hr at least 3 times sequentially to capture cell divisions and dying cell behaviours.
dfMT thicknesses (defined as the mean width of a given dfMT) was manually measured at high magnification using the Slice mode in Imaris and the Line tool for measuring the distance between two points. The innermost and outermost microtubule cable was defined as perimeter for the width measurements. Measurements were taken in 3 dfMT positions that covered the minimum, medium, and maximum width within a given dfMT to obtain an overall range distribution.
*EB3-GFP* tracking was done in Fiji using the MTrackJ plugin[3] to trace microtubules over time and create Kymographs, measure the velocity, track length, and comet duration. Tracking was also performed using the Spots and Surface tools in Imaris. The settings were initially automatically detected to fit the fluorescence and subsequently modified manually for improved fit. The settings were not changed between treatment groups.
Axon branch number quantifications were performed by drawing a 50µm line across the dorsal, medial, and ventral caudal fin edge region at a distance of 100µm parallel to the fin edge (which was determined to be the most reliable region to quantify distal branch numbers). Axons traversing this line were counted and averaged per treatment group. Axonal fluorescence measurements were taken using the Imaris Spots function by manually placing spots along the axon and using the fluorescence intensity profile measurements MatLab tool.

HyPer fluorescence was measured in the oxidized (505nm) and unoxidized (420nm) channels by using the region of interest (ROI) function in ZEN Black (Zeiss) whereby ROIs were selected inside highly magnified mitochondria (HyPer-mito), cytoplasm (HyPer-cyto) or the plasma membrane (HyPer-CAAX). The data was exported, and ratios determined in Excel. $H_2O_2$ measurements in zebrafish transiently injected with *tp63*:Hyper were manually performed by placing spot objects in Imaris 9.5.1 onto individual cells such that the spots extended to the lateral edges. Fluorescence intensities were measured inside the spots and 505/420nm ratios were subsequently calculated.

Membrane/cytoplasmic/nuclear ratio measurements for Nox1 staining we determined by measuring the fluorescence intensity in Fiji using single slice and single channel modes. First, a line using the line tool was placed along the plasma membrane that was clearly distinguishable from the cytoplasm due to increased fluorescence compared with the cytoplasm in all treatment groups. The measurement tool was subsequently selected to measure the mean fluorescence intensity. Three lines per caudal fin keratinocyte were averaged for 5 cells per animal and at least 3 animals per treatment group. To determine the mean cytoplasmic fluorescence intensity, nuclei were first overlaid with Nox1 fluorescence to ensure the exclusion of nuclear measurements. These images were then used to draw lines within the cytoplasm as above. For nuclear measurements, the line was placed inside a nucleus such that it spanned the longest extent of the nucleus. The fluorescence data was exported to Excel and the ratios calculated.

**Python Code to generate polar plots**

Polar plots were generated using Measurement points in Imaris to trace the X/Y coordinates (position) of microtubules. The positions were subsequently processed in Python to generate the polar plots using the following code:

```
import numpy as np
import matplotlib.pyplot as plt
import matplotlib as mpl
import sys, math
from locale import atof

def colorFader(c1,c2,mix=0): #fade (linear interpolate) from color c1 (at mix=0) to c2 (mix=1)
    c1=np.array(mpl.colors.to_rgb(c1))
    c2=np.array(mpl.colors.to_rgb(c2))
    return mpl.colors.to_hex((1-mix)*c1 + mix*c2)

# setting the axes projection as polar
ax = plt.axes(projection = 'polar')

S = 4
max = 0.0
with open(sys.argv[1]) as f:
    c = 0
    l = f.readline()
    while (l):
        if l[0] == ' ':
            c += 1
            S[c] = []
            l = (f.readline()).split()
            x_0 = atof(l[0])
            y_0 = atof(l[1])
```

```
            S[c].append((0.0, 0.0))
        else:
            S[c].append((atof(l[0]) - x_0, atof(l[1]) - y_0))
        l = (f.readline()).split()
c1='black'
c2='grey'

for x in S.keys():
    color = colorFader(c1,c2,x/c)
    X = list()
    Y = list()
    for rad in S[x]:
        x_1 = rad[0]
        y_1 = rad[1]
        if x_1 != 0:
            r = math.sqrt(x_1*x_1 + y_1*y_1)
            t = math.atan(y_1/x_1)
            if r > max:
                max = r

            X.append(t)
            Y.append(r)
            #plt.polar(t, r, color = 'black', linestyle='dashed', marker='k.', markersize=3)
        else:
            #plt.polar(0, 0, color = 'black', linestyle='dashed', marker='k.', markersize=3)
            X.append(0)
            Y.append(0)
    plt.polar(X, Y, 'k.', zorder=3, markersize=0, linestyle='solid', linewidth = 1.0, color = color)

#display the Polar plot
plt.show()
```

**Statistical analyses**

Comparisons of two groups were done using Student's t-test. Comparisons of more than two groups and a single variable were done using one-way ANOVA, whereas comparisons of multiple groups and multiple variables were done using two-way ANOVA. All comparisons were performed at alpha=0.05 (95% confidence interval) and post comparison tests, such as Tuckey's or Bonferroni's, as suggested by default in Prism 9. Significance is denoted as $p<0.05$, $p<0.01$, $p<0.001$, $p<0.0001$. The standard error of the mean (s.e.m.) is shown for all experiments given that at least two biological replicates were used.

**Mouse experiments**

Animal care and treatment

C57BL/6 mouse, 6 weeks old (15-22 g), were IP injected with 20 mg/kg paclitaxel (Athenex). The mice in each cage were randomly allocated to different treatment groups. Food and water were available *ad libitum* and experiments were performed during the light cycle (7:00 am to 7:00 pm). Animals were euthanized via $CO_2$ asphyxiation, followed by cervical dislocation. Any subjects that showed behavioral disturbances unrelated to chemotherapy-induced pain were excluded from further behavioral testing. The animals were breed and care in the DVR core facility (University of Miami) according to established NIH protocols and handled in strict agreement with good animal practice as approved by the appropriate Institutional Animal Care and Use Committee (IACUC).

Tissue staining
At the end point (48h) the paws and the skin at the back were collected, placed in 10% formalin, and stored overnight at 4°C. The samples were embedded in paraffin and sectioned at 7µm. Sections were deparaffinized, washed with PBS, and incubated at room temperature for 5 min in 0.1% Triton X-100 in PBS and then 30 min in blocking solution (5% BSA). Sections were incubated with the primary antibody diluted in blocking solution, overnight at 4°C in a humidity chamber. Following PBS washes, sections were incubated for 1 h at room temperature with a secondary antibody. ProLong reagent (Invitrogen) was used for mounting. Antibodies used were against detyrosinated tubulin Millipore AB3201, p-KIF11 Invitrogen PA5-38647, alpha-tubulin Proteintech 66031-1-Ig, mouse Alexa Fluor 488 Invitrogen A21202, and rabbit Alexa flour 546 Invitrogen A10040. DAPI was used for nuclei counterstaining.

Imaging
Immunofluorescence staining were observed with widefield microscopy using a Plan-Apochromatix 100× objective lens (oil immersion, numerical aperture [NA] 1.4), 20x and 10x objective lens on inverted Zeiss Axio Observer Z1 using AxioVision 4.8 software. Images were acquired using a monochrome Zeiss Axio Cam MRm CCD camera. Images were lastly processed with the NIH Image J software.

Mouse RNAseq analysis: The RNAseq data set was published in Cirrincione et al. Data (2022)[5]. Following differential gene expression analysis (FDR<0.01) comparing paclitaxel-treated to vehicle control animals (n=4), identified genes were queried in gProfiler and grouped by their molecular function. Using normalized gene counts, *Kif* genes were analysed for their expression and plotted over time. Significance values comparing paclitaxel and vehicle controls were subsequently established using Prism 9 (GraphPad). *Kif11* was queried in STRING to identify associated networks. Interacting proteins were compared to the RNAseq data set and differentially expressed genes were graphed in Prism 9.

**Human RNAseq**

All studies were approved by the Mayo Clinic's IRB review board. Three female breast cancer survivors were recruited who received standard adjuvant paclitaxel treatment. Paclitaxel was administered in these patients over the course of 12 wees by infusions of 80 mg/m$^2$ each. A retrospective review of the patient's medical records was conducted to confirm the date of CIPN diagnosis (5 weeks for XMMP003, 31 weeks for XMMP002, and 35 weeks for XMMP001 prior to the performed skin biopsies) and record the administration of paclitaxel to the patient. Neurological history, examination, and QLQ-CIPN20 quality of life questionnaire, which has been validated to detect the presence of CIPN[6], was completed in all subjects. Consent was given to obtain two skin punch biopsies of approximately 4 mm diameter. These were collected 10 cm proximal to the lateral malleolus on the distal leg as described previously[7]. The biopsies were carried out under anesthesia following local injection of 2% lidocaine with epinephrine, using a sterile technique. In addition, three healthy volunteers of similar age and same gender (female) were recruited as controls. Volunteers serving as controls had no history of neuropathy, diabetes, or familial neuropathies. One skin biopsy was used for RNA sequencing (Azenta/Genewiz) whereas the other biopsy was used elsewhere. RNA isolation and quality control were performed, and paired-end Illumina sequencing (HiSeq 2x150bp) was conducted. Sequence reads were trimmed to remove possible adapter sequences and nucleotides with poor quality using Trimmomatic v.0.36. The trimmed reads were mapped to the Homo sapiens GRCh38 reference genome available on ENSEMBL using the STAR aligner v.2.5.2b. The STAR aligner is a splice aligner that detects splice junctions and incorporates them to help align the entire read sequences. BAM files were generated through this step. Unique gene hit counts were calculated by using featureCounts from the Subread package v.1.5.2. The hit counts were summarized and reported using the gene_id feature in the annotation file. Only unique reads that fell within exon regions were counted. If a strand-specific library preparation was performed, the reads were strand-specifically counted. After extraction of gene hit counts, the gene hit counts table was used for downstream differential expression analysis. Using DESeq2, a comparison of gene expression between the customer-defined groups of samples was performed. The Wald test was used to generate p-values and log2 fold changes. Genes with an adjusted p-value < 0.05 and absolute log2 fold change > 1 were called as differentially expressed genes for each comparison. Below are the results of the number of significantly differentially expressed genes for all comparisons provided. A gene ontology analysis was performed on the statistically significant set of genes by implementing the software GeneSCF v.1.1-p2. The goa_human GO list was used to cluster the set of genes based on their biological processes and determine their statistical significance. A list of genes clustered based on their gene ontologies was generated.

Significantly differentially expressed candidate genes for control subjects and CIPN patients were subsequently graphed either as normalized gene counts or relative expression using GraphPad Prism 9. Upregulated KIF genes were queried in STRING to identify networks. The data was uploaded to the Gene Omnibus Database with the GEO Accession number GSE228633.

**ZStretcher User Manual**

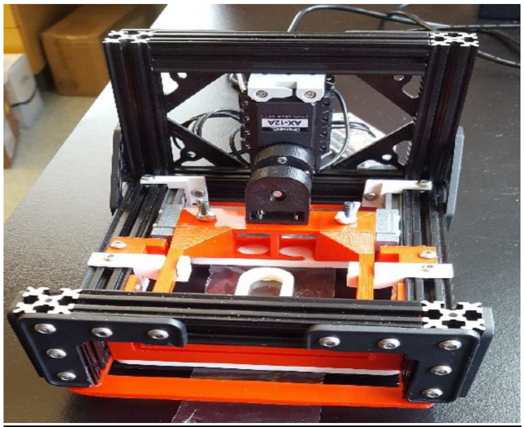

Contents
Overview
Components & Materials
Device Set-Up
LabVIEW Operation
Notes

**Overview**
This is the fourth version of the zebrafish stretcher device and software. With most of the components being3D printed, the platform can be reprinted to match various stage inserts for confocal microscope, but the version below is shown for common Zeiss confocal microscope stages. The LabVIEW program that runs the motor must be installed on whichever device will be used with the controller (either the microscope's desktop computer or a laptop computer). There should be no need to access the block diagram of the LabVIEW program, as modifications to the block diagram will alter the calibrations of the motor operation.

<u>Components & Materials</u>
Film & Mount
Film Blueridge Films, Inc. BFI-1880 Metallocene
Silicone grease and embedding agarose gel
3D PLA dividers: zfish chamber – _horizontal/vertical/double partition.stl
Frame
Beams and Joints from OpenBeamUSA.com
Constructed by previous teams at UMaine
Bolts tightened with 5/64 hex/allen key
Sliding Platform
3D PLA prints attached to metal sliding brackets
Retained from the 3rd version of the model
Original manual caliper saved
Bolt Housing
3D PLA print:motor base.stl
(1) M5-.8 T-Nut and (4) M2-.4 screws
Fasteners
(2) 5mm tall 3D PLA prints
(2) 2mm tall 3D PLA print: fasteners.stl
(4) Original hex bolts and wing nuts
Stepper Motor
Electronics
Dynamixel AX-12A and Servo Manager Kit from Trossen Robotics
U2D2 Controller, 12V 5A power supply, AX/MX power hub 200mm 3 pin cables, power jack adapter, USB M-F cable
Hardware
(1) 3D PLA adapter: motor attachment.stl
(1) M5-.8 25mm bolt, (1) M5-.8 T-Nut, (4) M2-.4 12mm bolts
(1) Dynamixel base, (2) OpenBeamUSA bolts, (4) M2-.4 5mm bolts

**Device Set-Up**
The stretcher features a stationary clamp and a sliding clamp which is moved by a linear actuator to stretch a Metallocene film. The Dynamixel servo has a head attachment, which allows the bolt to be attached. Currently, a 25mm bolt is attached, but the attachment can be unscrewed and the 50mm bolt can be swapped into place if needed. The bolt has a 0.5mm diameter. The stepper motor, bolt, and bolt housing are all permanently attached to the device, so no set-up is needed.

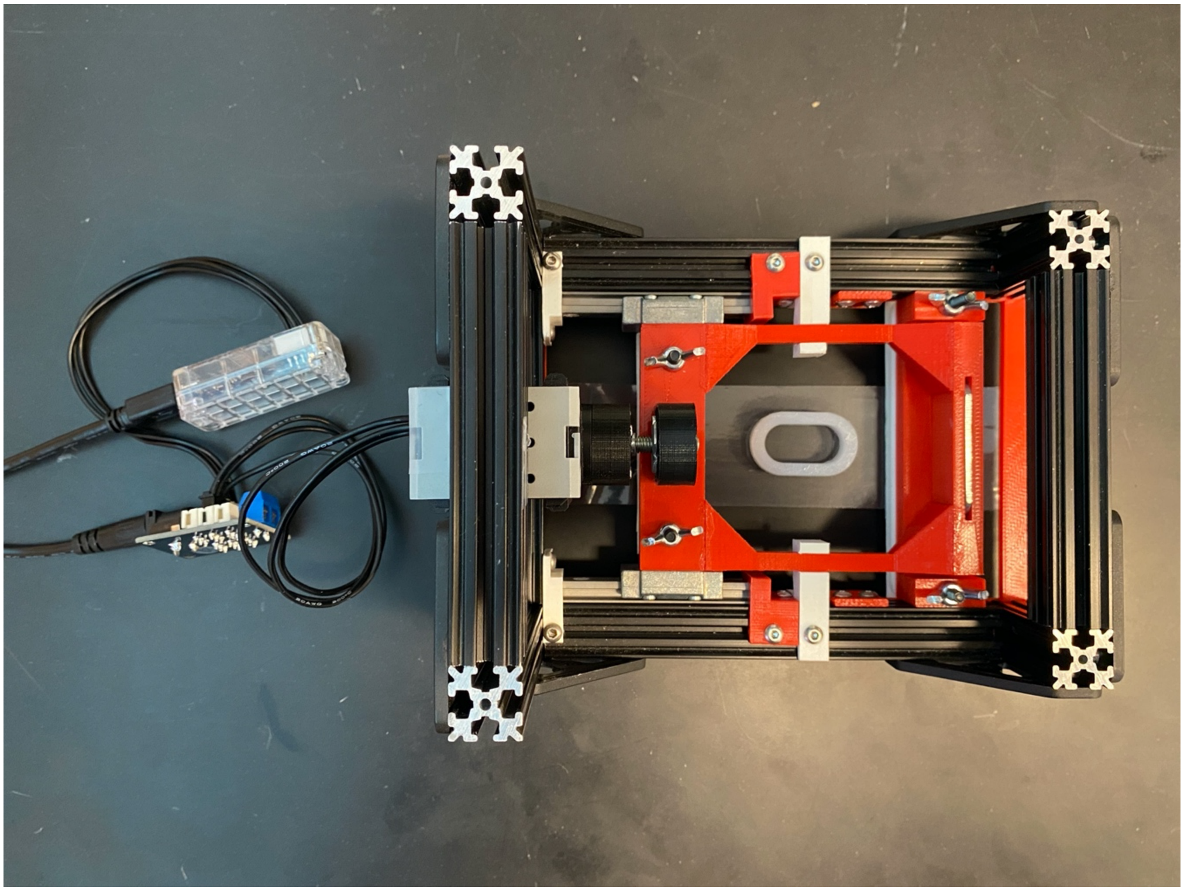

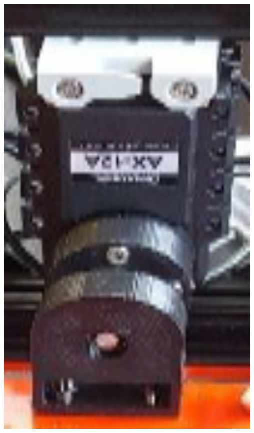

The film is fixed in place by plastic holders, and manually tightened by wing nuts and bolts. To insert the film between the white and black fasteners, the bolts from the bottom must be loosened. To stay within the working distance of the Zeiss Plan Neofluar 20X/0.50 Infinity objective, the black fasteners closest to the objective must be kept in place. However, do not tighten the fasteners entirely until the stepper motor is reset for maximal stretching.

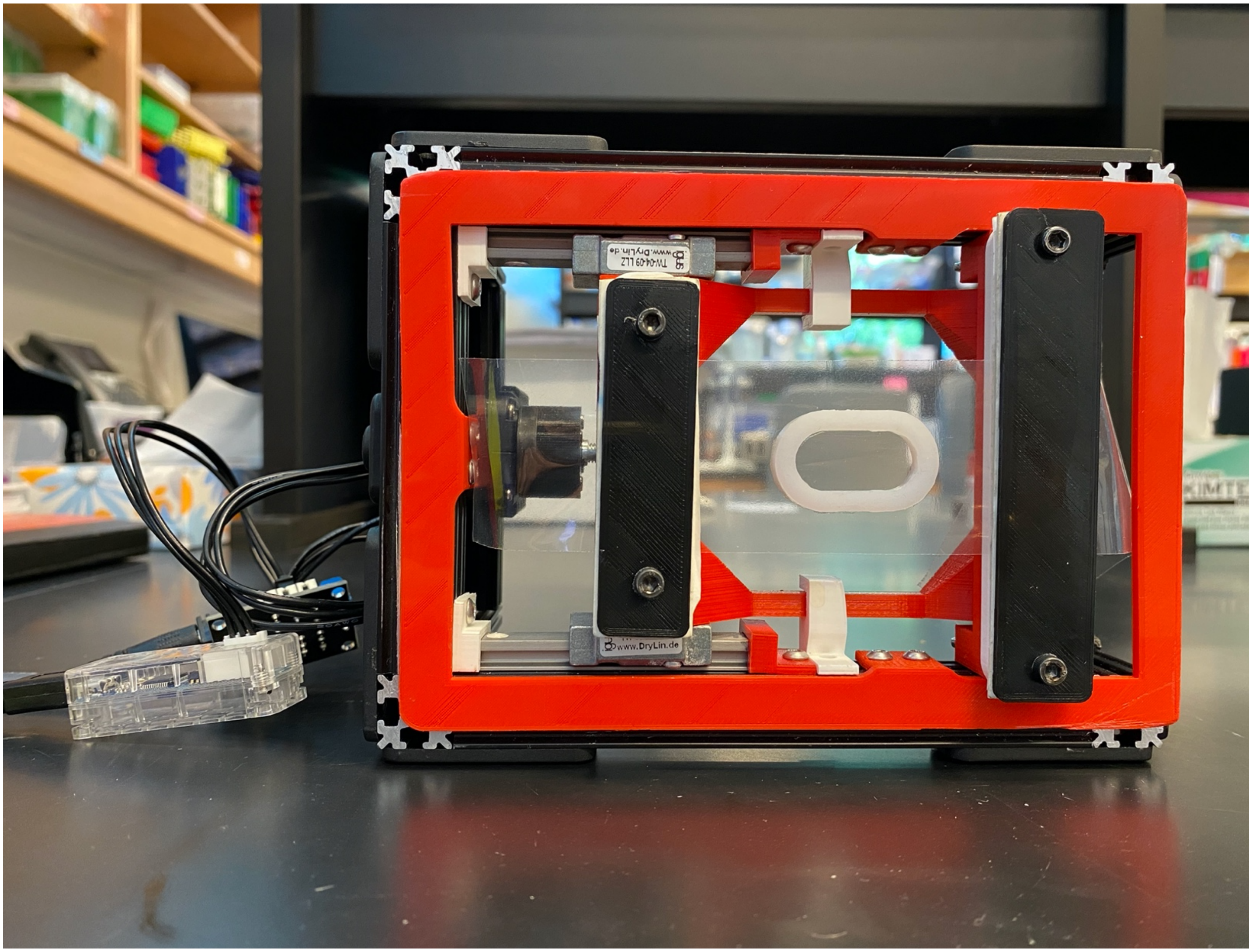


The stepper motor is controlled by a controller, a power hub, and a power supply. The black/black 3pin cable should be connected to the motor power hub as seen below. The U2D2 controller should be plugged into the power hub via the TTL port with the black 3pin cable seen below. The microUSB end of the USB cable should be inserted into the U2D2 controller, and the regular end of the USB cable should be inserted into the computer that is used for operation. The power supply should be plugged into a regular power outlet and into the power hub.

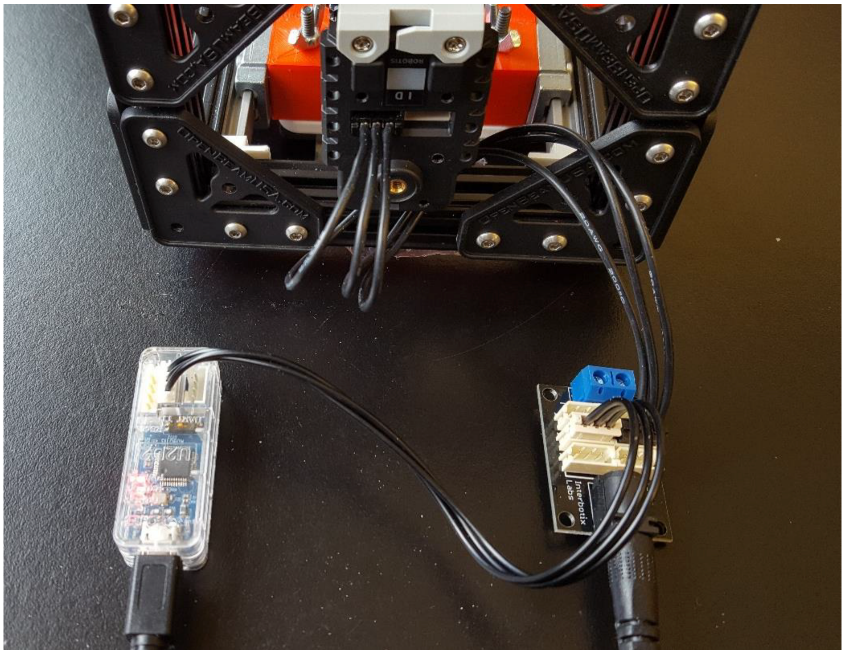
Interbotix
Labs

**LabVIEW Operation**
The current virtual instrumentation file (.vi) is the ZStretcher_June18.vi (supplied by Rieger lab upon request), which also requires the Dynamixel Setup sub.vi to operate. The Dynamixel library may not be included in the program upon startup, so the “dynamixel library” folder holds the .dll and .h files used by LabVIEW. It can be imported by selecting “Tools” → “Import” → “Shared Library” (use this link for directions: LabVIEW). Upon opening the ZStretcher_June18.vi, the window below will appear. Annotated are the 9 essential tabs.

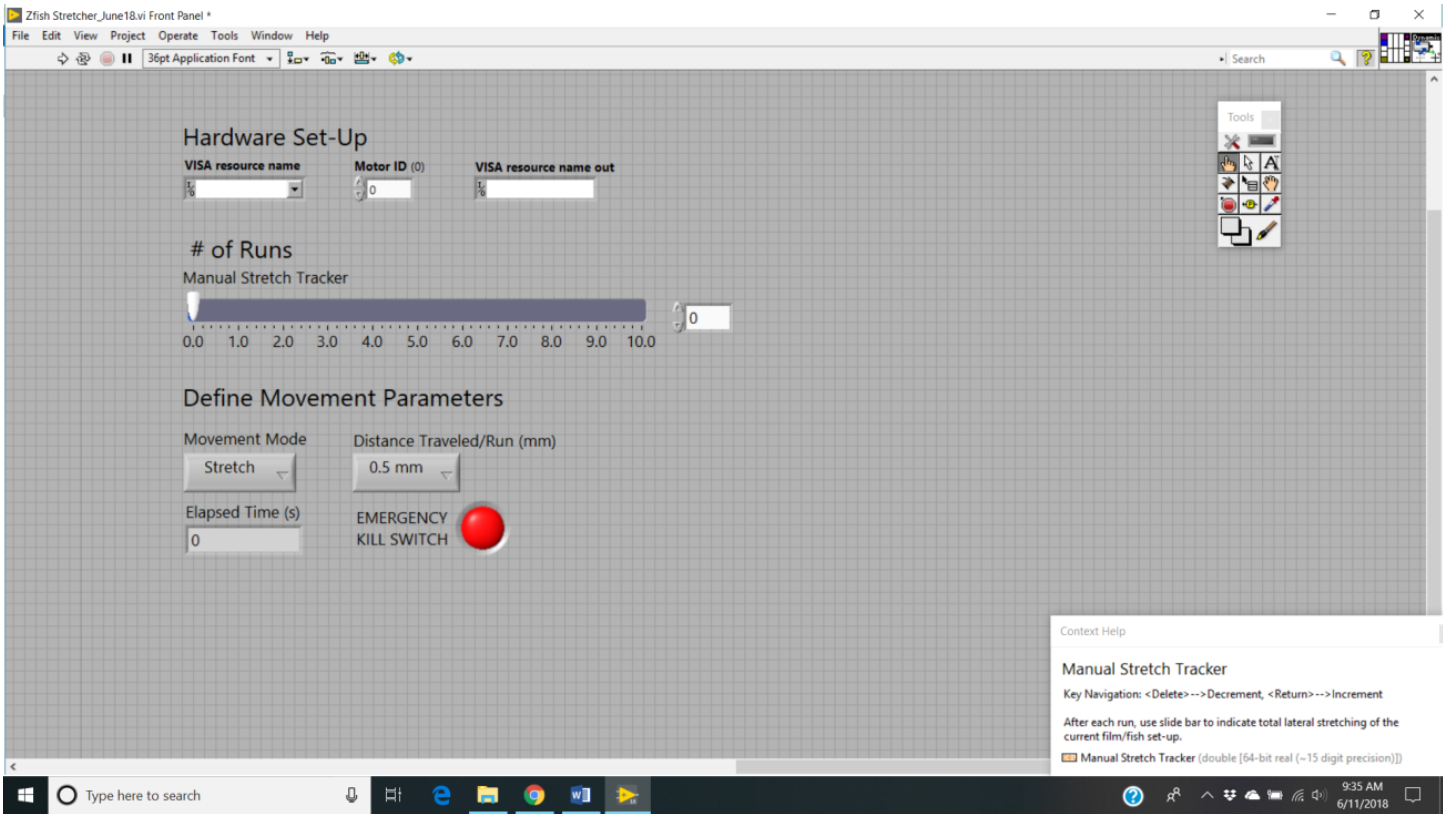


1. The arrow executes the program as defined on what is called the front panel
   a. The front panel is simply the window shown.
   b. Operations 6 & 7 define the operations that occur during a run/execution

2. The red circle is what ends each program, and must be selected at the end of each run
   a. Clicking this button before the completion of the run/execution will result in a short circuit which causes the program to never stop
      i. Should this happen, simply unplug the power sources and restart the program

3. The VISA resource name corresponds to the port on the computer which the controller, via the USB is plugged into
   1. This can be checked once the USB is plugged into your computer and accessing the device
      manager
   b. Once in the device manager, click on “Ports” _and see what port appears with the USB Serial Port

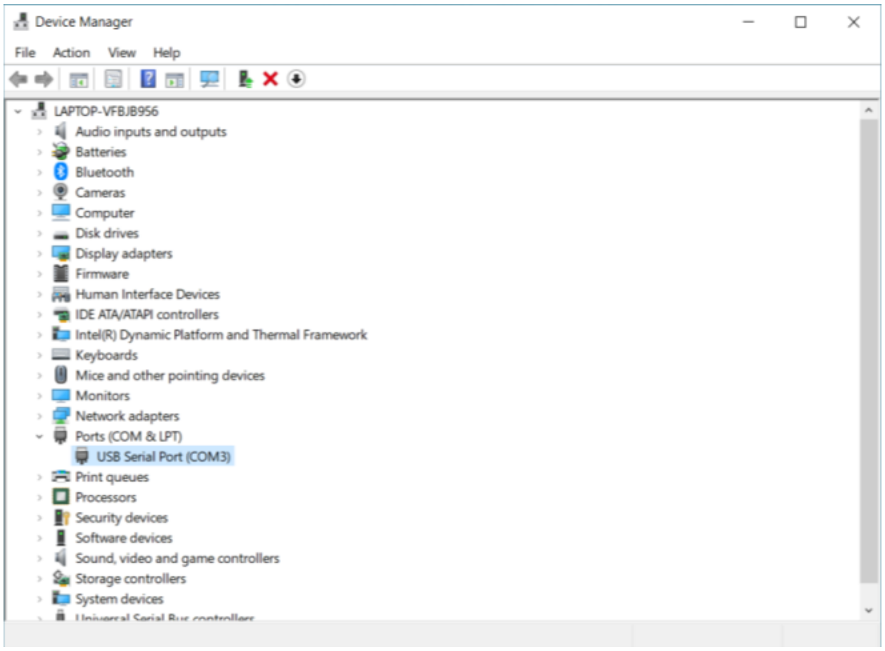


4. The Motor ID is what the .vi is currently calling the Dynamixel AX-12A.

   a. The default is 0

      i. If the program never runs, open the control panel via clicking "Window" → "Show Block Diagram"

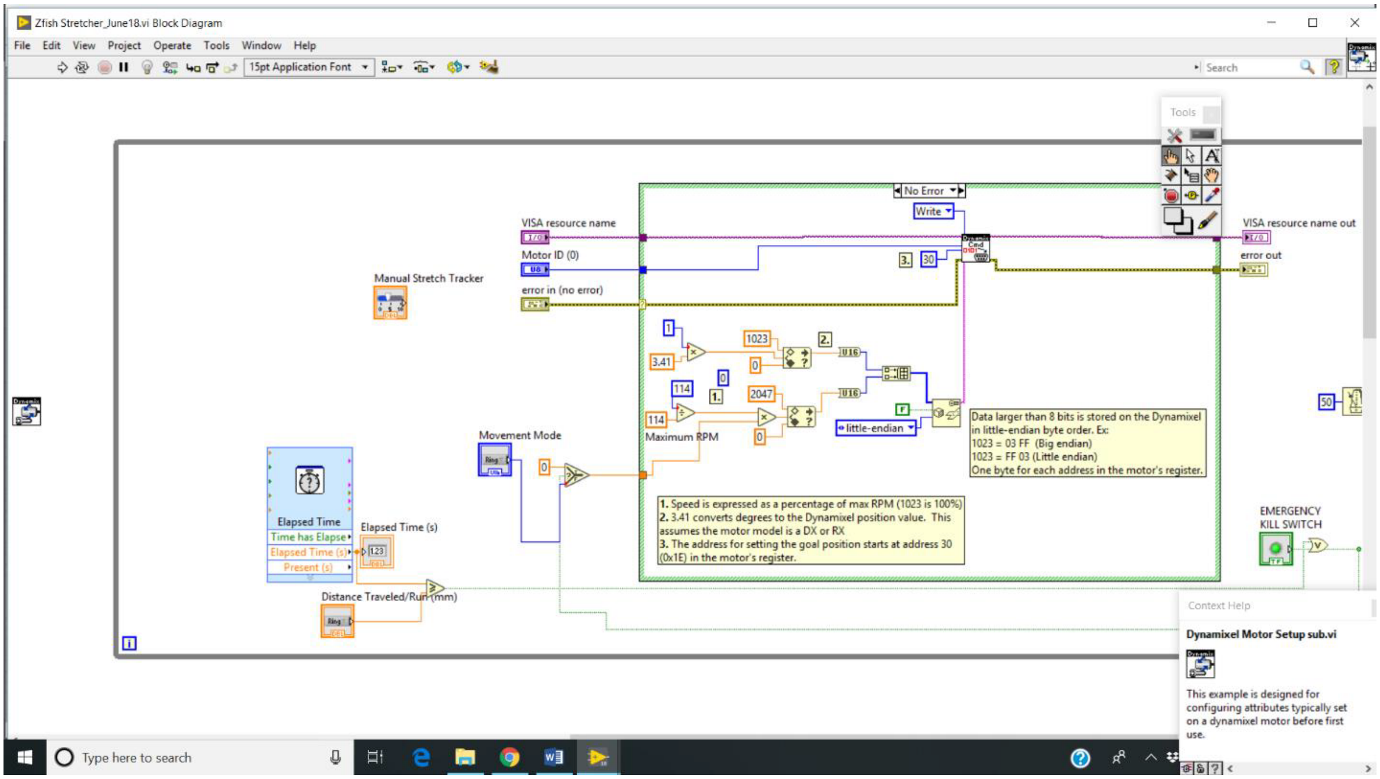

ii. Click on the box indicated, which is the sub.vi

iii. Check to make sure it is calling the motor 0, or manually correct

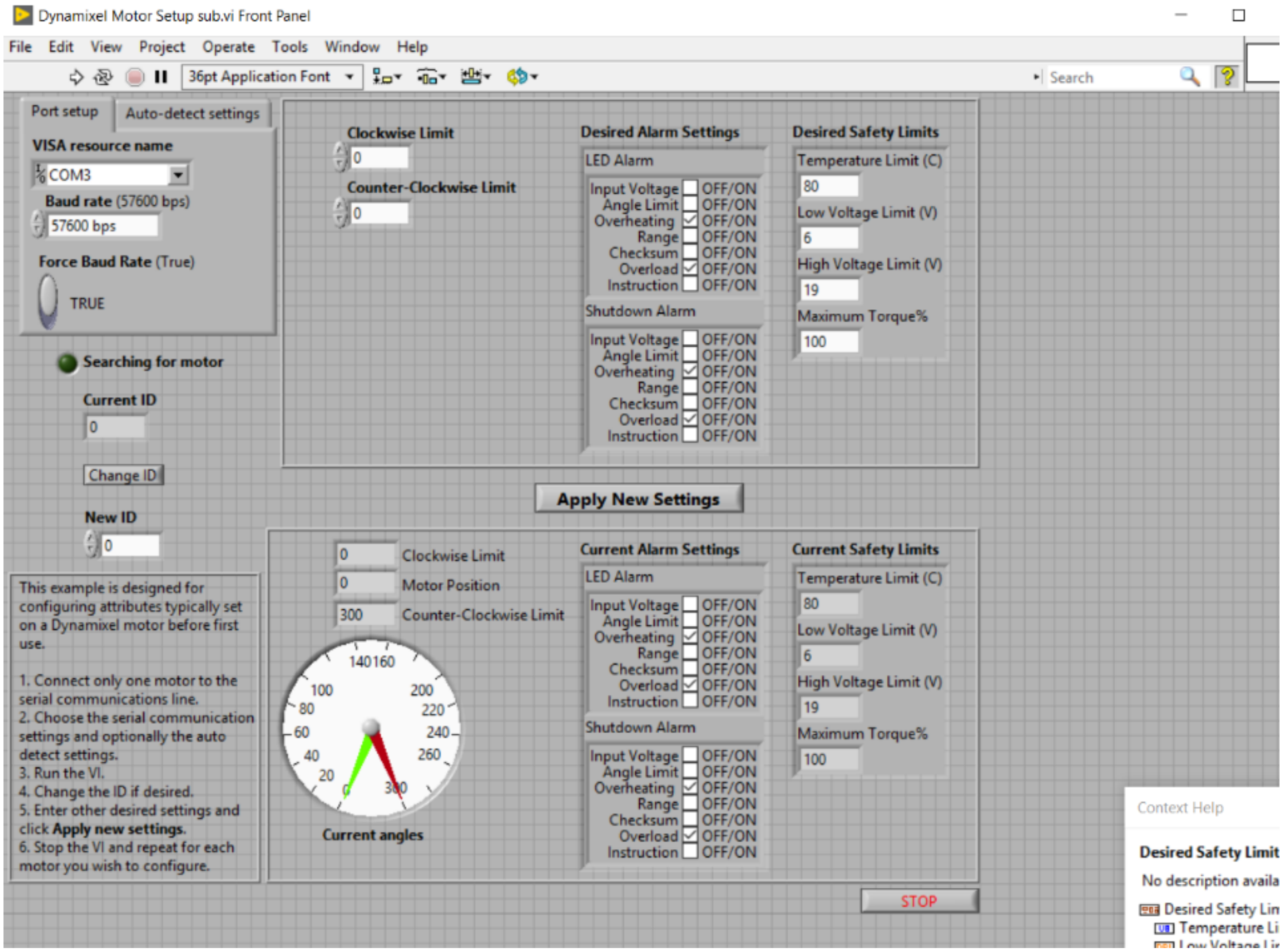


iv. After changing “New ID” _to 0 and running the program, be sure to save
v. Exit the sub.vi and return to the main .vi

5. This is a manual indicator of how many runs/executions have been done on the current sample
   a. Hit enter, or click on the up indicator, after each program run
   b. At the end, you can simply multiply the # of runs by the number in box 7 to get the total distance travelled

6. Movement Mode is a click box to either stretch the fish or to reset the device to have the bolt moved back
   a. Multiple runs may be needed to return the bolt to an optimal start position

7. The Distance Travelled/Run is a dropdown menu that includes 0.5/1.0/0.4/0.8/Reset options
   a. Reset should be used with the Reset Movement Mode
   b. It is advised that once you are in stretching mode, to only use one distance

8. Elapsed time indicates how long the program runs each time
a. Distance travelled is controlled by the RPM and time i. Each distance is set at the same RPM, but
the time changes
ii. Ideal times are respectively 6.25/11.4/5.3/9.4/15 seconds

9. The Emergency Kill Switch can be clicked during the program to cease the run

You can scroll over the controls on the front panel and a description of each will appear in the bottom right text box on the screen. Before each experiment, it is wise to reset the bolt back so that it is barely in the housing/nut. This may need to be executed multiple times.

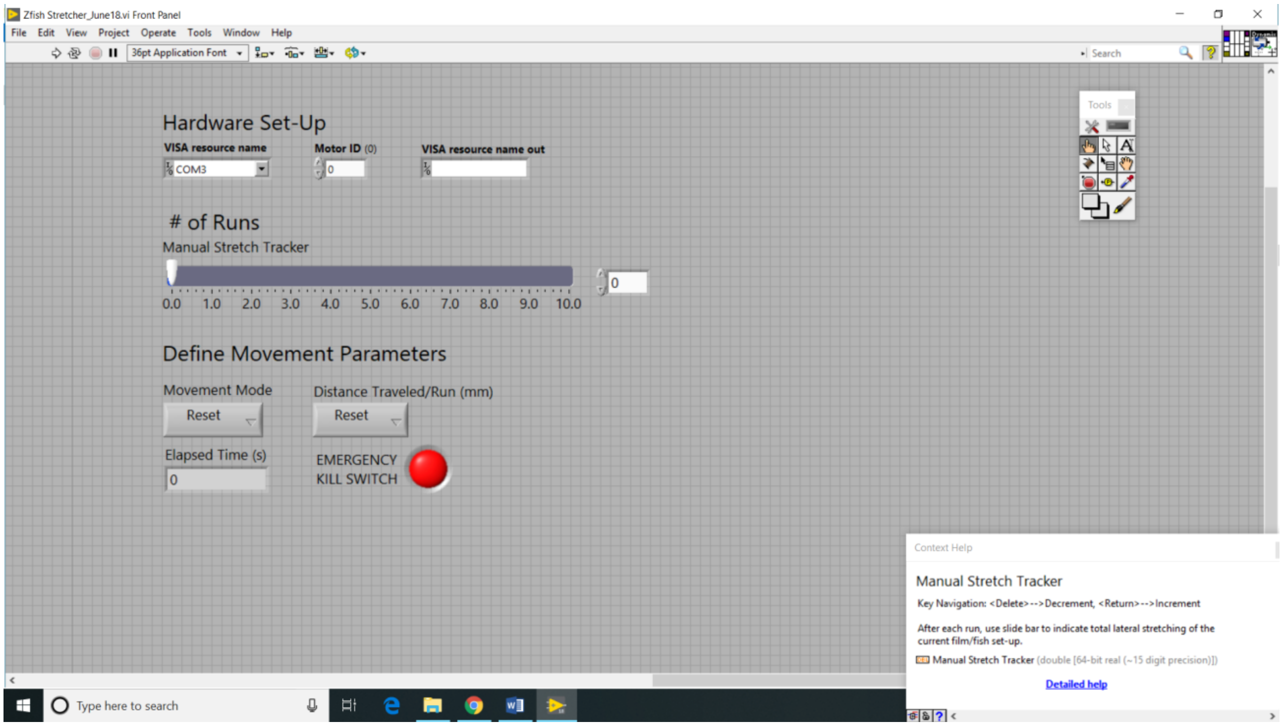


After the bolt is reset, you can then run the program on stretch mode and define your distance used per iteration. Be sure to mark the # of runs at the end of each execution.

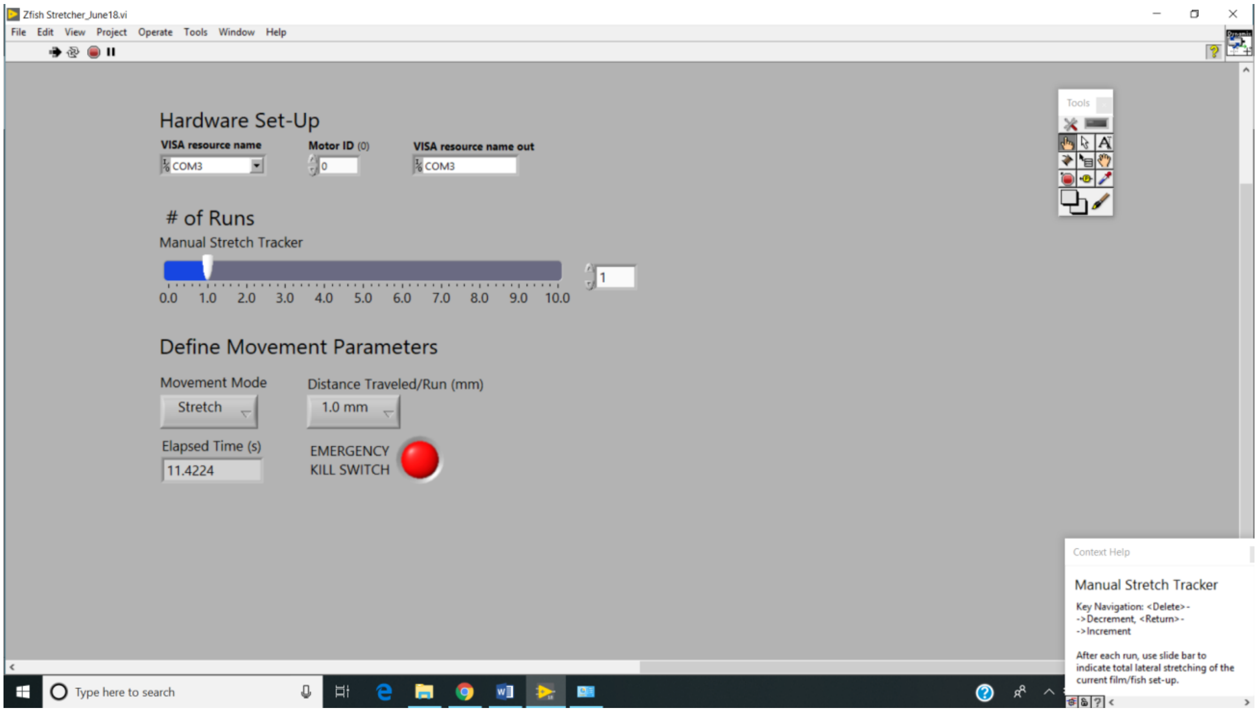


This LabVIEW program has a bug that cannot be fixed, and which may depend on the version. A warning pop-up may appear (shown below) during the middle of your run that states that the control blocks cannot find the angle of the motor during the run. Since we use time to control distance, this error is irrelevant. Simply click on “Continue” and the run will remain normal. However, if you click “Stop” or in rare instances, the motor will continue to spin, try the kill switch or unplug the power sources and restart the program.

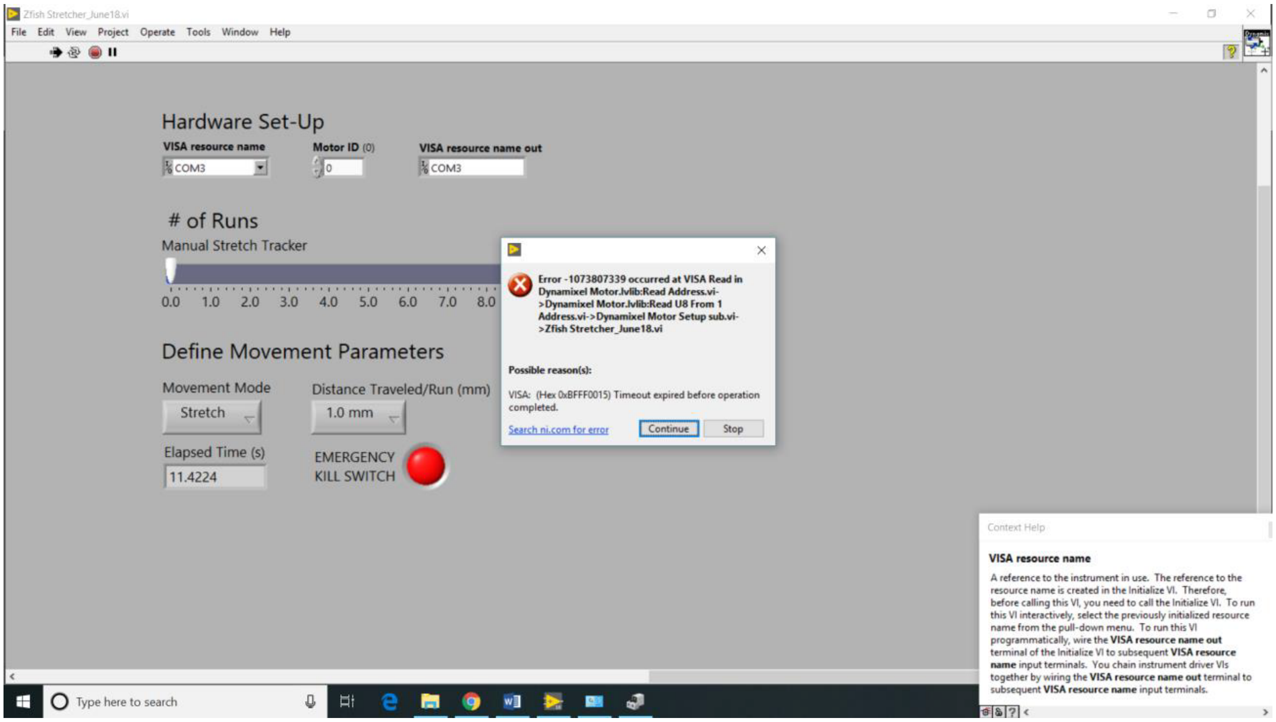

**Notes**

1. LabVIEW must be installed on whatever device is being used to operate the device.
2. Viewing the block diagrams is great to understand the of the program but be careful to not save unwanted edits.
3. View the user manual for the Dynamixel AX-12A.
4. Websites for Trossen Robotics and Metallocene BFI 1880 (Blueridge Film).
5. All hardware was purchased from Home Depot, printed with an UltiMaker 2
6. To periodically check the accuracy of the travel distance, run the 0.4mm or 0.8mm multiple times
    - The 0.4mm option should result in a ½ rotation of the motor and the 0.8mm should result in one full rotation of the motor
    - If adjustments are needed, right click on the Distance Travelled box and selecting “Date Entry”, then “Edit Items” will allow you to adjust the time corresponding to each distance.

**Supplemental References**


1 Lisse, T. S. *et al.* Paclitaxel-induced epithelial damage and ectopic MMP-13 expression promotes neurotoxicity in zebrafish. *Proceedings of the National Academy of Sciences of the United States of America* **113**, E2189-2198, doi:10.1073/pnas.1525096113 (2016).

2 O'Brien, G. S. *et al.* Coordinate development of skin cells and cutaneous sensory axons in zebrafish. *Journal of Comparative Neurology* **520**, 816-831, doi:10.1002/cne.22791 (2012).

3 Meijering, E., Dzyubachyk, O. & Smal, I. Methods for cell and particle tracking. *Methods Enzymol* **504**, 183-200, doi:10.1016/B978-0-12-391857-4.00009-4 (2012).

4 Nieuwenhuis, J. & Brummelkamp, T. R. The Tubulin Detyrosination Cycle: Function and Enzymes. *Trends Cell Biol* **29**, 80-92, doi:10.1016/j.tcb.2018.08.003 (2019).

5 Cirrincione, A., Reimonn, C., Harrison, B. & Rieger, S. Longitudinal RNA sequencing of skin and DRG neurons in mice with paclitaxel-induced peripheral neuropathy. *Data* **7**, 72, doi:https://doi.org/10.3390/data7060072 (2022).

6 Postma, T. J. *et al.* The development of an EORTC quality of life questionnaire to assess chemotherapy-induced peripheral neuropathy: the QLQ-CIPN20. *Eur J Cancer* **41**, 1135-1139, doi:10.1016/j.ejca.2005.02.012 (2005).

7 Engelstad, J. K. *et al.* Epidermal nerve fibers: confidence intervals and continuous measures with nerve conduction. *Neurology* **79**, 2187-2193, doi:10.1212/WNL.0b013e3182759608 (2012).
